# Analysis of altered pre-mRNA splicing patterns caused by a mutation in the RNA binding protein hnRNPA1 linked to amyotrophic lateral sclerosis

**DOI:** 10.1101/2022.02.03.479052

**Authors:** Yeon J. Lee, Donald C. Rio

## Abstract

Amyotrophic lateral sclerosis (ALS) is a debilitating neurodegenerative disease characterized by loss of motor neurons. Human genetic studies have linked mutations in RNA binding proteins as causative for this disease. The hnRNPA1 protein, a known pre-mRNA splicing factor, is mutated in a number of ALS patients. Here, we generate two cell models to investigate how a mutation in the C-terminal low complexity domain of hnRNPA1 affects global pre-mRNA splicing patterns and RNA binding. We show that a single amino acid change in the C-terminal low complexity domain (D262V) leads to changes in splicing of thousands of transcripts whose genes are linked to the DNA damage response, cilia organization and translation. We also show that there are changes in RNA binding of the mutant hnRNPA1 protein to transcripts whose splicing patterns change. Finally, we show that cells expressing the hnRNPA1 D262V mutation exhibit an aggregation phenotype, markedly reduced growth rates and changes in stress granules. This study shows that global changes in pre-mRNA splicing patterns caused by a single mutation in the hnRNPA1 protein lead to phenotypes related to ALS and that specific cellular pathways are affected.

## Introduction

Amyotrophic lateral sclerosis or ALS is the third most common adult onset neurodegenerative disorder worldwide. It is generally characterized by progressive paralysis starting at the limbs and ultimately leading to death caused by respiratory failure. There is no cure and current treatments fail to slow the progression of the disease. It is known that mutations in RNA binding proteins (RBPs), such as FUS, TIA1 and TDP43 are linked genetically to ALS (Kwiatkowski et al. 2009; Vance et al. 2009; Kim et al. 2013; Johnson et al. 2014; Mackenzie et al. 2017). In addition, mutations in heterogeneous nuclear ribonucleoproteins (hnRNPs), e.g. hnRNPA1 and hnRNPA2/B1, have been linked to familial cases of ALS (Kim et al. 2013; Liu et al. 2016; Beijer et al. 2021). Also, repeat expansions in the C9orf72 gene cause ALS and are associated with the formation of RNA foci containing low solubility hnRNP proteins suggesting that RNA toxicity via the sequestration of RBPs plays a role in the disease (Cooper-Knock et al. 2014; Conlon and Manley 2017; Balendra and Isaacs 2018).

Furthermore, hnRNPA1 and A2/B1 are often found sequestered in cytosolic stress granules together with other RBPs causative for ALS, such as FUS and TDP-43. Mutations identified in the prion-like low complexity domains of hnRNPA1 and A2/B1 are found in subsets of ALS patients (Kim et al. 2013; Liu et al. 2016; Beijer et al. 2021). Under normal conditions, these hnRNP proteins have an intrinsic tendency to self-aggregate because of their prion-like domains, but this tendency can often be abnormally increased because of the disease-causing mutations (Kim et al. 2013; Geuens et al. 2016).

The RNA binding splicing factor hnRNPA1 contains two RRM-type RNA binding domains at the N-terminus and glycine-arginine-rich and prion-like low complexity domains at the C-terminus. Single point mutations within this C-terminal domain have been associated with neurodegenerative diseases, such as ALS. There are 4 known ALS-associated mutations found in the C-terminal domain of hnRNPA1 (Liu et al. 2016; Beijer et al. 2021). The aspartic acid (D) to valine (V) mutation at position 262 is the only disease-associated variant that impacted an evolutionally conserved residue in ALS patients (Kim et al. 2013). Hence, we focused our efforts on testing the effects of this mutation in hnRNPA1 using genome editing and expression of this mutant protein on global pre-mRNA splicing patterns, target RNA binding and phenotypic characterization in two human cell line models. Either expression of hnRNPA1 D262V in 293 HEK cells or editing the D262V mutation into the endogenous locus in neuronal SH-SY5Y cells led to marked changes in the patterns of RNA splicing of thousands of transcripts, some linked to the DNA damage response and cilium organization. In some cases, the splicing pattern changes could be linked to alterations in binding of the hnRNPA1 protein using *in vivo* irCLIP RNA binding assays. Interestingly, either expression of the hnRNPA1 D262V mutant protein from either an inducible cDNA or from the edited endogenous locus caused the cells to clump and aggregate, resulted in a marked reduction in cell growth rate and changes in stress granules. This behavior might explain, in part, the neuronal cell death caused by this mutation in ALS patients. This study provides further mechanistic insights into how familial mutations in the hnRNPA1 gene might contribute to the ALS disease phenotype.

## Results

### Expression of the hnRNPA1 D262V mutant protein in HEK 293 cells and characterization of global RNA pre-mRNA splicing patterns and *in vivo* RNA binding

To test the effects of the hnRNPA1 D262V mutation, our initial approach was to generate inducible cell lines to express either hnRNPA1 wild type or D262V mutant epitope-tagged cDNAs. These cells were generated using FLP-recombinase mediated integration in 293 cells, while expressing endogenous wild type hnRNPA1. This experimental set up was initially designed to mimic ALS patients carrying a heterozygous hnRNPA1 D262V mutation. FLP-In 293 cells expressing hnRNPA1 wild type or D262V mutant epitope-tagged cDNAs were induced with tetracycline for 72 hr. Cells were harvested and immunoblot analysis was performed to confirm the induction of the exogenous epitope-tagged hnRNPA1 proteins (Fig. 1A). Interestingly, we observed that the expression of the endogenous hnRNPA1 protein decreased slightly upon expression of the exogenous hnRNPA1 wild type or D262V mutant proteins (Fig.1A). We also observed that hnRNPA1 D262V mutant protein expression is slightly lower than hnRNPA1 wild type expression when normalized to endogenous beta-actin protein as a loading control.

**Figure 1.**
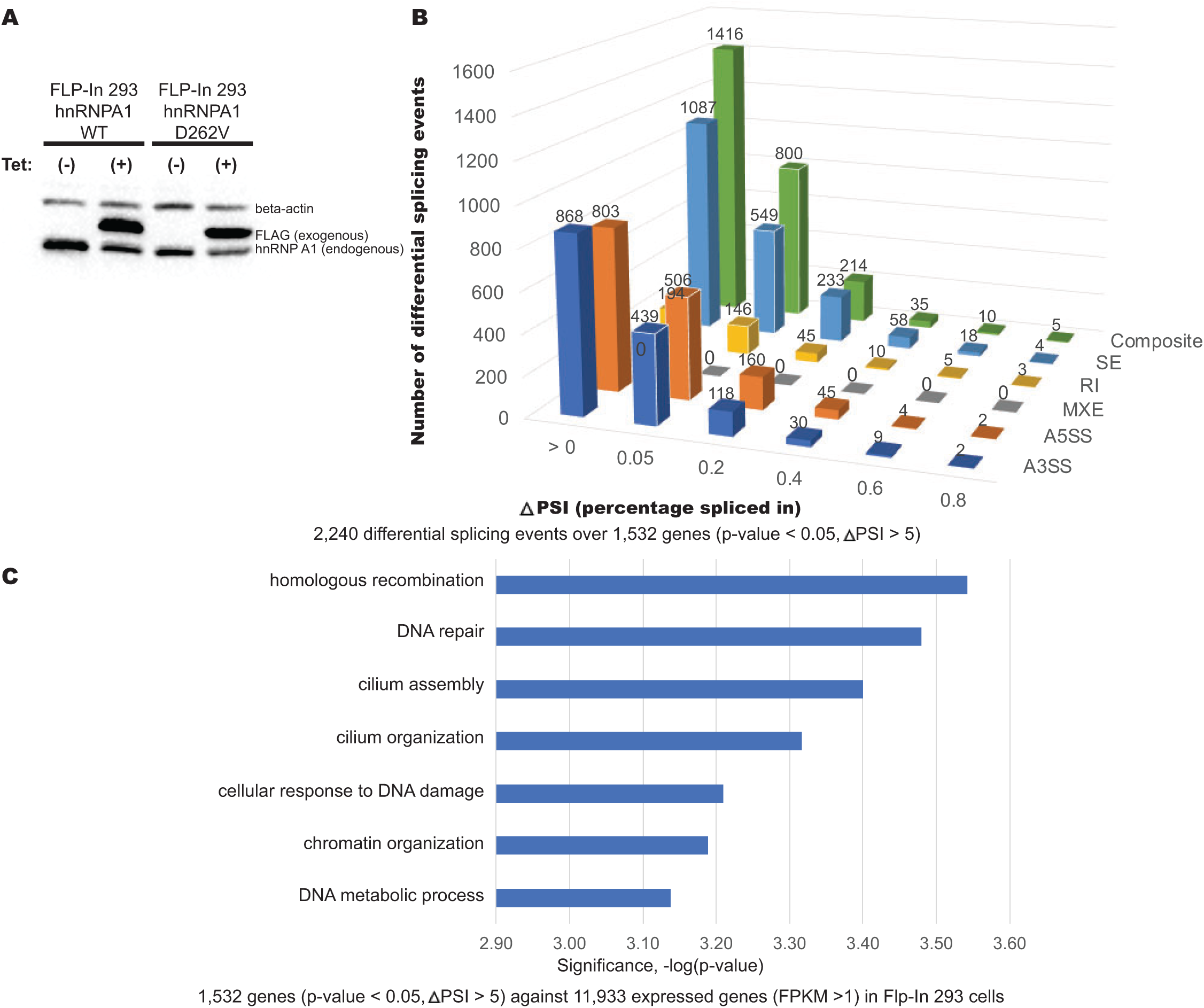
Quantitation of differential alternative splicing events controlled by hnRNPA1 D262V mutant. **A**. Tet-responsive induction of FLP-In™ T-Rex™ 293 cells expressing WT hnRNPA1 and ALS-associated D262 mutant. Tetracycline (Tet)-inducible 293 FLP-In hnRNPA1 wild type and hnRNPA1 mutant (D262V) cells were induced with 1 μg/mL Tetracycline (Tet) for 72 hr prior to harvest. Protein lysates were immunblotted with hnRNPA1 endogenous hnRNPA1), FLAG (exogenous hnRNPA1), and beta-actin serves as loading controls. **B**. Quantitation of differential alternative splicing events controlled by hnRNPA1 D262V mutant. JUM (Junction Usage Model or JUM) is a splicing-annotation independent method for determining pre-mRNA splicing patterns from RNA-seq data. Only splice junction-spanning reads are counted for quantitation. This resulted in quantitative comparison of AS events (2,240) whose splicing patterns were significantly altered upon expression of FLP-In 293 hnRNPA1 D262V samples versus the control (p < 0.05, ΔPSI >5), covering 1532 genes. **C**. A graph of Gene Ontology (GO) term enrichment of hnRNPA1 D262V mutant splicing targets from JUM analysis in FLP-In 293 hnRNPA1 D262V cells.

To examine the effect of the hnRNPA1 D262V disease-causing mutation on pre-mRNA splicing, we generated RNA-seq libraries from the FLP-In 293 cells expressing either epitope-tagged hnRNPA1 wild type or mutant proteins. These libraries, prepared in triplicate, were sequenced to approximately 35-50M PE100 reads each. After mapping of the reads to the human genome (hg38), quantitative alternative splicing (AS) analysis was performed using JUM software (Wang and Rio 2018) to detect differential splicing (DS) events between the wild type and D262V mutant cells. JUM (Junction Usage Model) is a splicing-annotation independent computational method for quantitating and comparing pre-mRNA splicing patterns from RNA-seq datasets (Wang and Rio 2018). JUM only uses splice junction-spanning reads for splicing isoform quantitation and delineates standard and “composite” splicing patterns. This computational analysis resulted in a quantitative comparison of AS events (2,440) whose splicing patterns were significantly altered upon expression of Tet-inducible FLP-In hnRNPA1 D262V samples versus the wild type hnRNPA1 control (p < 0.05, △ PSI >5), covering 1,532 genes (Fig. 1B). All categories of splicing events, with an exception of mutually exclusive exons (MXE), seem to be affected by expression of the hnRNPA1 D262V mutant. Gene Ontology (GO) enrichment analysis of high-confidence hnRNPA1 D262V mutant splicing targets from the JUM analysis against 11,933 expressed genes (FPKM > 1) in 293 cells revealed these differentially spliced targets are involved in homologous recombination, DNA repair, cilium assembly, cellular response to DNA damage stimulus and chromatin organization (Fig. 1C). This analysis suggests that altered splicing patterns caused by expression of the hnRNPA1 D262V mutant may lead to cellular DNA damage stress, which could explain the growth defects observed with these cells (see below). Differential expression analysis comparing FLP-In 293 hnRNPA1 D262V to the wild type revealed only 345 genes are differently expressed by DEseq2 analysis (FDR p < 0.05, FC > 1.5) (Suppl. Fig. S1). These genes are involved in guanine metabolic process, protein targeting to ER, protein localization to membrane, and SRP-dependent co-translational protein targeting to membrane (Suppl. Fig. S1).

### Effects of expressing the hnRNPA1 D262V mutant from the endogenous locus in genome-edited human SH-SY5Y neuronal cells

In order to test the effects of expression of the hnRNPA1 D262V mutant protein in human neuronal cells, we used genome editing at the endogenous hnRNPA1 locus in SH-SY5Y, human neuroblastoma cells. This approach allowed us to generate homozygous hnRNPA1 mutant cell lines using Cas9-RNP transfection. DNA sequencing of genomic PCR-amplified DNA from isolated edited single cell clones confirmed that the edited SH-SY5Y hnRNPA1 D262V mutant cell lines are homozygous for the mutation. We generated RNA-seq libraries from SH-SY5Y cells expressing either the unedited control hnRNPA1 locus and the genome edited hnRNPA1 D262V mutant genes. These libraries, prepared in triplicate, were sequenced to approximately 70-130M PE100 reads each and were mapped to the human genome (hg38). We performed JUM bioinformatic analysis to determine changes in pre-mRNA splicing patterns comparing the edited cells carrying the hnRNPA1 D262V mutation with the unedited control cells. This analysis resulted in a quantitative comparison of AS events (4,368) whose splicing patterns were significantly altered upon expression of the hnRNPA1 mutant D262V protein versus the wild type control (p < 0.05, △ PSI >5), covering 3,322 genes (Fig. 2A). All categories of splicing patterns were affected. GO term enrichment analysis of 3,322 differentially-spliced (DS) transcripts against 11,394 expressed genes (FPKM >1) in SH-SY5Y cells indicated that genes with altered splicing patterns were involved in cellular metabolic processes, cilium organization, chromatin remodeling, DNA repair, and cytoskeleton organization (Fig. 2B). The GO term analysis suggests that cellular metabolic or DNA damage stress might result from the splicing alterations caused by expression of the hnRNPA1 D262V mutant protein. Differential expression analysis comparing SH-SY5Y hnRNPA1 D262V to unedited control cells revealed that only 619 genes are differently expressed using DEseq2 analysis (FDR p < 0.05, FC > 1.5) (Suppl. Fig. S2). The GO-terminology of differentially expressed genes indicates these genes are involved in centrosome localization, microtubule organizing center localization and gene silencing by miRNA (Suppl. Fig. S2).

**Figure 2.**
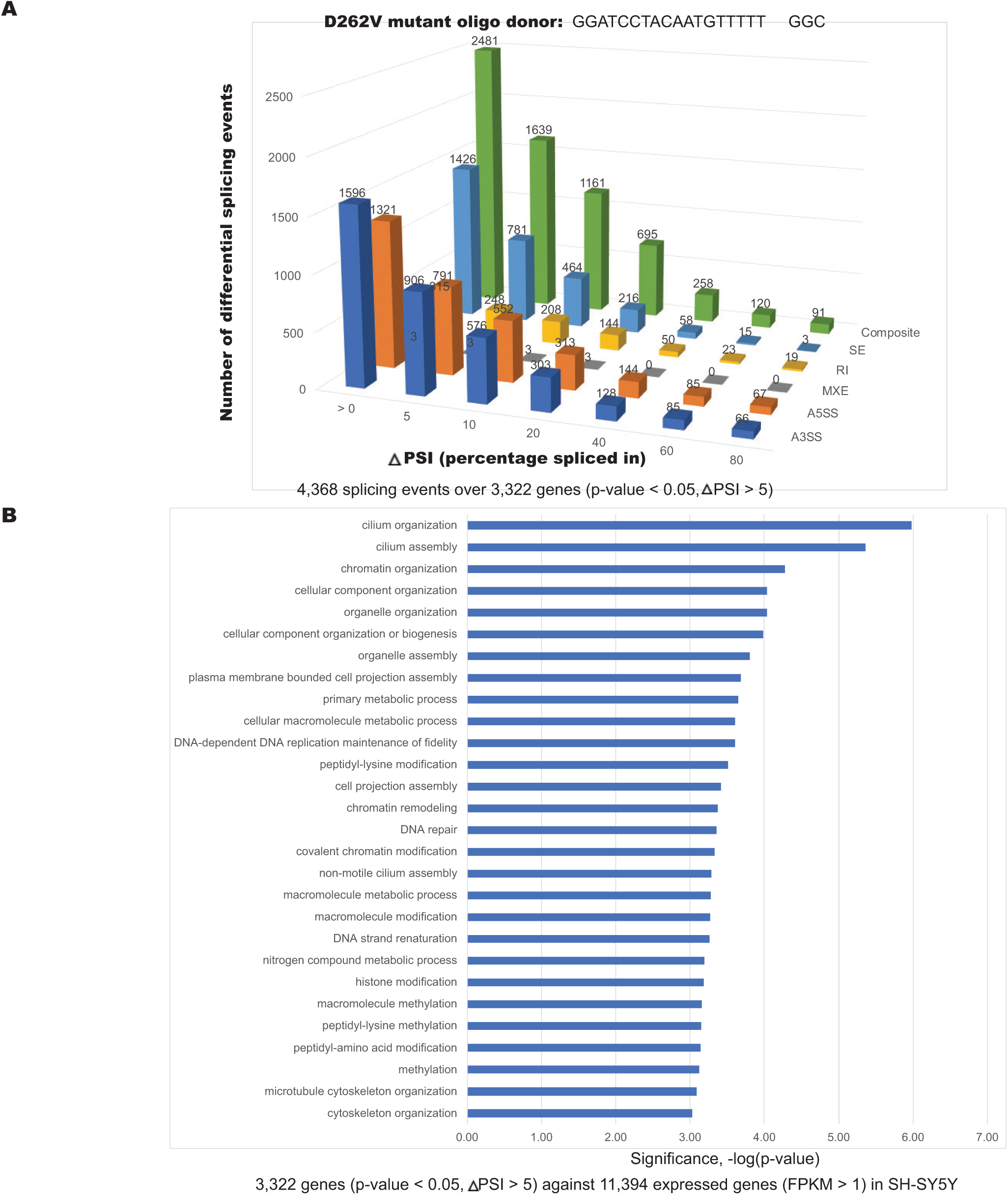
Over 4,000 differential splicing events were detected in genome-edited SH-SY5Y neuroblastoma cells expressing ALS-associated hnRNPA1 D262V mutation. **A**. Differential AS events detected upon expression of ALS-associated hnRNPA1 D262V mutation. Over 4,000 splicing events were significantly altered in hnRNPA1 D262V samples in neuronal cell line versus the control (with p < 0.05, ΔPSI >5). Shown on the y-axis is the number of differential splicing events and difference in percentage spliced in (or PSI) shown on the x-axis. The splicing events were filtered based on the delta PSI. **B**. A graph of Gene Ontology (GO) term enrichment of differentially spliced transcripts in hnRNPA1 D262V mutant protein-expressing SH-SY5Y cells.

We compared DS events in FLP-In 293 cells and SH-SY5Y cells expressing the ALS-associated hnRNPA1 D262V mutant protein. A Venn diagram shows the overlapping DS targets resulting from the two different cell lines expressing the hnRNPA1 D262V mutant resulted in 639 high-confidence splicing targets that were detected by JUM analysis (Fig. 3A). These high-confidence splicing targets are involved in DNA repair, cellular response to DNA damage stimulus, cellular metabolic processes, chromatin remodeling, and cilium organization and assembly (Fig. 3B). It is interesting that GO enrichment analysis of high-confidence DS events in two different cell line models expressing the hnRNPA1 D262V mutant resulted in differentially spliced transcripts from genes involved in DNA repair, cilium assembly and chromatin organization, cellular pathways that have been in implicated in neurodegeneration diseases (Chen et al. 2013; Madabhushi et al. 2014; Hardiman et al. 2017).

**Figure 3.**
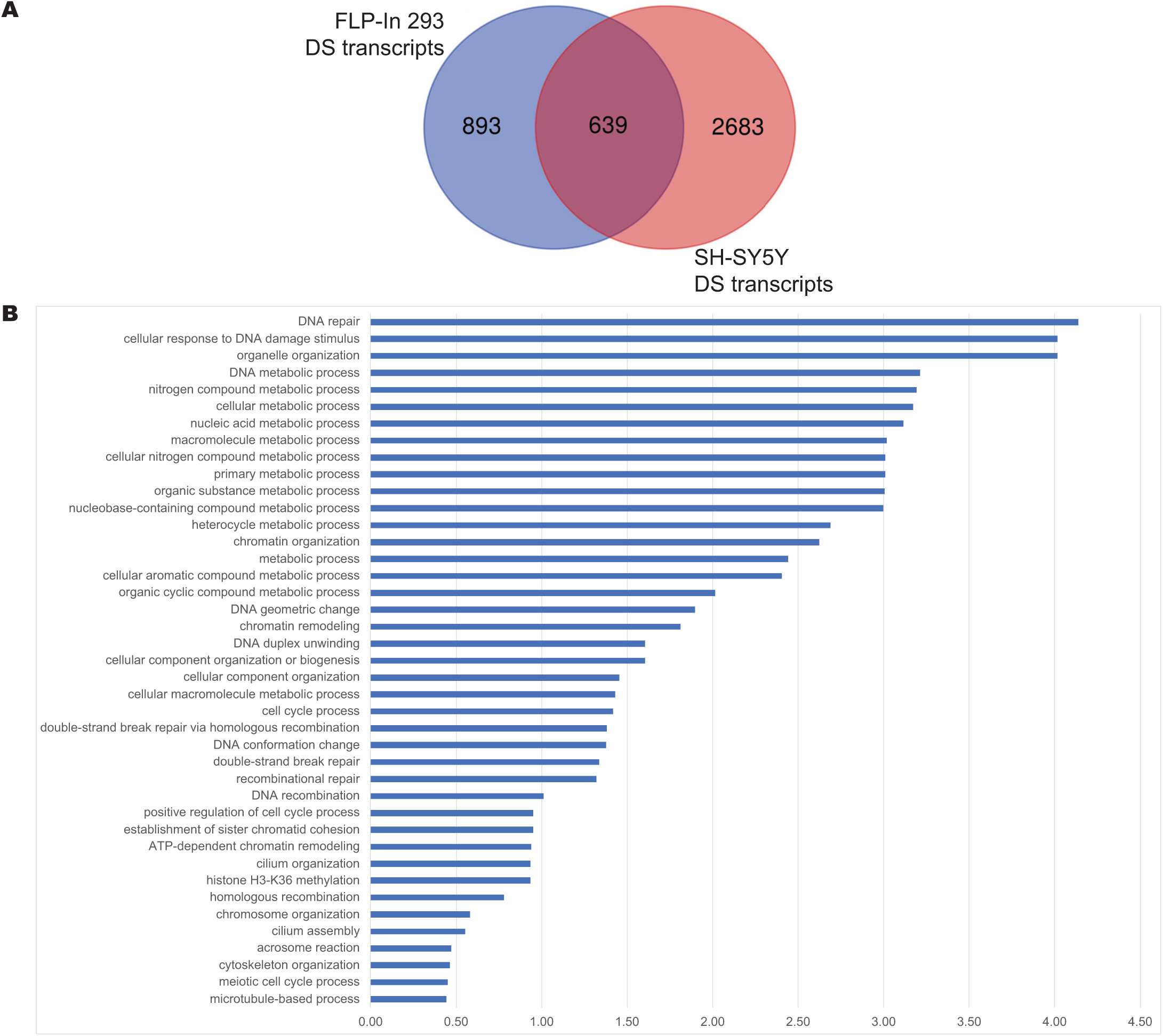
Comparison of differentially spliced (DS) transcripts in FLP-In 293 cells and SH-SY5Y expressing ALS-associated hnRNPA1 D262V mutation. **A**. Venn diagram showing the overlapping splicing targets resulting from 2 different cell lines expressing hnRNPA1 mutant proteins, resulted in 639 high-confidence splicing targets that were detected by JUM. **B**. A graph of Gene ontology (GO)-term enrichment analysis of hnRNPA1 D262V mutant splicing targets was performed against all expressed genes in FLP-In 293 cells and in SH-SY5Y neuroblastoma cells. Enriched GO term categories included response to DNA damage stimulus, DNA repair and cilium assembly that includes cytoskeletal proteins.

### Expression of the hnRNPA1 D262V mutant in either FLP-In 293 or SH-SY5Y cells causes cell aggregation and a markedly reduced growth rate

Interestingly, we observed a cell aggregation phenotype in hnRNPA1 mutant cell lines in both in FLP-In 293 and SH-SY5Y cells compared to wild type, as well as a markedly slower growth rate (Suppl. Fig. S3). FLP-In 293 cells expressing hnRNPA1 wild type and the D262V mutant were induced and immunofluorescence assays (IFA) were performed using anti-FLAG antibody to detect exogenously expressed hnRNPA1 wild type and D262V mutant proteins (Suppl. Fig. S3A). Remarkably, cells expressing the hnRNPA1 D262V mutant protein displayed a distinct phenotype in which the cells aggregate in the FLP-In 293 cells. We also noted that the D262V mutant expressing cells had a slower growth rate compared to the wild type cells. We treated FLP-In 293 cells with tetracycline for 1, 3, 5, and 7 days and then counted viable cells. The growth curves shown (Suppl. Fig. S3B) clearly display a growth delay in cells expressing the hnRNPA1 D262V mutant protein compared to wild type-expressing cells.

A similar phenotype was observed in genome-edited SH-SY5Y cells expressing the hnRNPA1 D262V mutant protein. An equal number of SH-SY5Y cells were plated and cultured for 10 days followed by crystal violet staining. There was a dramatic difference in cell density between SH-SY5Y cells expressing either the hnRNPA1 wild type or D262V mutant proteins, again displaying a slow growth rate in the D262V mutant cells compared to wild type (Suppl. Fig. S3C). Interestingly, the cell aggregation phenotype observed in the 293 cells was also observed in

SH-SY5Y cells expressing the hnRNPA1 D262V mutant protein compared to the wild type. Differential interference contrast (DIC) microscopic images of SH-SY5Y cells expressing hnRNPA1 wild or D262V mutant proteins show the cell aggregation phenotype of the mutant cells (Suppl. Fig. S3D). Taken together these data show that expression of the hnRNPA1 D262V mutant leads to changes in cell-cell contacts and a marked reduction in growth rate.

### The hnRNPA1 D262V mutant cells exhibit defects in stress granules

Recent studies suggest that the response of RNA metabolism to stress plays an important role in neurodegenerative diseases (Nussbacher et al. 2019; Wolozin and Ivanov 2019; Dudman and Qi 2020). RNA-binding proteins (RBPs) control the utilization of mRNAs during stress, in part through formation of stress granules (SG). SG are supposed to be transient structures, but the chronic stresses associated with aging lead to chronic and persistent SG that appear to initiate and foster the aggregation of disease-related proteins (Wolozin and Ivanov 2019; Marcelo et al. 2021).

To examine whether there is a difference in stress-granule formation under stress in hnRNPA1 262V mutant-expressing cells, we treated the neuronal cell line expressing either wild type or the D262V mutant with arsenite for 30 min., then followed SG assembly and disassembly by immunofluorescence using an eIF3α antibody. EIF3α has been shown to translocate to SG under stress and is an SG marker. hnRNPA1 staining is indicated in green and eIF3α staining is indicated in red (Suppl. Fig. S4). In order to observe if there is any difference in SG disassembly between hnRNPA1 wild type or D262V mutant cell lines, we also followed SG disassembly 2 hr. after removal of arsenite. We showed that not only that hnRNPA1 translocates to the cytoplasm under stress (indicated by higher blue staining in nucleus under arsenite treatment), but that there is a delay in SG disassembly in the hnRNPA1 D262V mutant cell line, indicating a possible difference in RNA metabolism. Notably, a recent study showed that two ALS-associated hnRNPA1 mutants, P288A and D262V, attenuate both SG formation and exhibited a delay in SG disassembly in a HeLa cell over-expression model (Beijer et al. 2021). Our findings also suggest that genome-edited SH-SY5Y cells expressing the hnRNPA1 D262V mutant protein exhibit a delay in SG disassembly compared to unedited control cells.

We also identified many transcripts encoding stress granule-associated proteins that were differentially spliced both in neuroblastoma SH-SY2Y and Tet-inducible 293 cells (Suppl. Fig. S5), including transcripts encoding RNA binding proteins that have been implicated in neurodegenerative disease and proteins shown to associate with other RBPs causative for ALS, such as FUS and TDP-43. These differentially spliced transcripts are encoded by genes whose products were associated with neuronal RNP granules, eIF4A, and SG (Suppl. Fig. S5).

### Analysis and comparison of *in vivo* RNA binding by wild type and D262V hnRNPA1 using irCLIP

The expression of the hnRNPA1 D262V mutant protein resulted in alternative splicing changes in thousands of transcripts. We wanted to determine if the hnRNPA1 D262V mutant protein might bind to a different subset of transcripts compared to the wild type hnRNPA1 protein. A common method for probing RNA-protein interactions *in vivo* involves UV cross-linking and immunoprecipitation (or CLIP) assays, which can be combined with high-throughput sequencing to generate transcriptome-wide binding maps of RNA-binding proteins (Darnell 2010; Hafner et al. 2010; Huppertz et al. 2014; Van Nostrand et al. 2016; Zarnegar et al. 2016). RNA binding proteins in covalent complexes with their bound RNAs from UV-irradiated cells are subjected to immunoprecipitation followed by RNase treatment. Protein-RNA adducts corresponding to comigrating species are recovered following SDS-PAGE, protein-associated RNA fragments are isolated and RT-PCR amplified followed by Illumina sequencing. We utilized one of the latest CLIP methods, called Quick irCLIP that incorporates an infrared-biotinylated adapter for the detection of protein-RNA interactions (Kaczynski et al. 2019). We performed irCLIP experiments in both inducible FLP-In 293 cells expressing hnRNPA1 wild type or D262V mutant epitope-tagged proteins and using SH-SY2Y cells endogenously expressing hnRNPA1 wild type or the D262V edited mutant.

We first optimized immunoprecipitation conditions for the hnRNPA1 protein from extracts derived from each cell line. Immunoblot analysis showed an efficient pulldown of the protein of interest (Suppl. Fig. S6A). Infrared scans of the nitrocellulose-bound immunoprecipitates using FLAG-antibody against exogenously expressed epitope-tagged hnRNPA1 wild type and D262V mutant in FLP-In 293 cells are shown indicating efficient ligation of the biotinylated adapter to the RNA in the hnRNPA1 immunoprecipitates (Suppl. Fig. S6B). The blue bar on the SDS-PAGE gel indicates the size-selected regions that were isolated for downstream RNA isolation and processing. For the SH-SY5Y cell lines, an anti-hnRNPA1 antibody was used to immunoprecipitate hnRNPA1 from the unedited wild type or endogenously edited hnRNPA1 D262V mutant cell line extracts (data not shown).

After removing PCR-duplicates, read mapping and peak calling, we detected over 480,000-580,000 clusters with at least 10 overlapping sequence reads after FDR filtering (FDR, p-value < 0.05) for each sample, which mapped to ∼8,000 genes (Fig. 4A). Next, we compared the differentially spliced target RNAs from our RNA-seq analyses to the transcriptome binding sites of the hnRNPA1 wild type and D262V mutant proteins in the neuronal cell lines. Approximately 65% of the differentially spliced transcripts were bound by either the hnRNPA1 wild type or D262V mutant proteins (Fig. 4B). These differentially spliced transcripts correspond to only ∼40% of the hnRNPA1 binding target RNAs determined by irCLIP. This observation may not be surprising since besides its role in pre-mRNA splicing, hnRNPA1 is also known to be involved in transcriptional regulation, mRNA transport, translation, miRNA biogenesis and telomere maintenance (Jean-Philippe et al. 2013; Clarke et al. 2021). Searching the irCLIP binding clusters for enriched motifs found similar logos that resembled the hnRNPA1 *in vitro* SELEX binding motif (Burd and Dreyfuss 1994) and were observed in both cell lines expressing wild type and D262V mutant hnRNPA1 (Fig. 4C). This observation may not be too surprising since the disease-associated hnRNPA1 mutation resides within the low-complexity domain and not within RNA binding RRM domains. There is biochemical evidence that the C-terminal low complexity domain of hnRNPA1 mediates protein-protein interactions (Cartegni et al. 1996; Jean-Philippe et al. 2013; Wang and Rio 2018).

**Figure 4.**
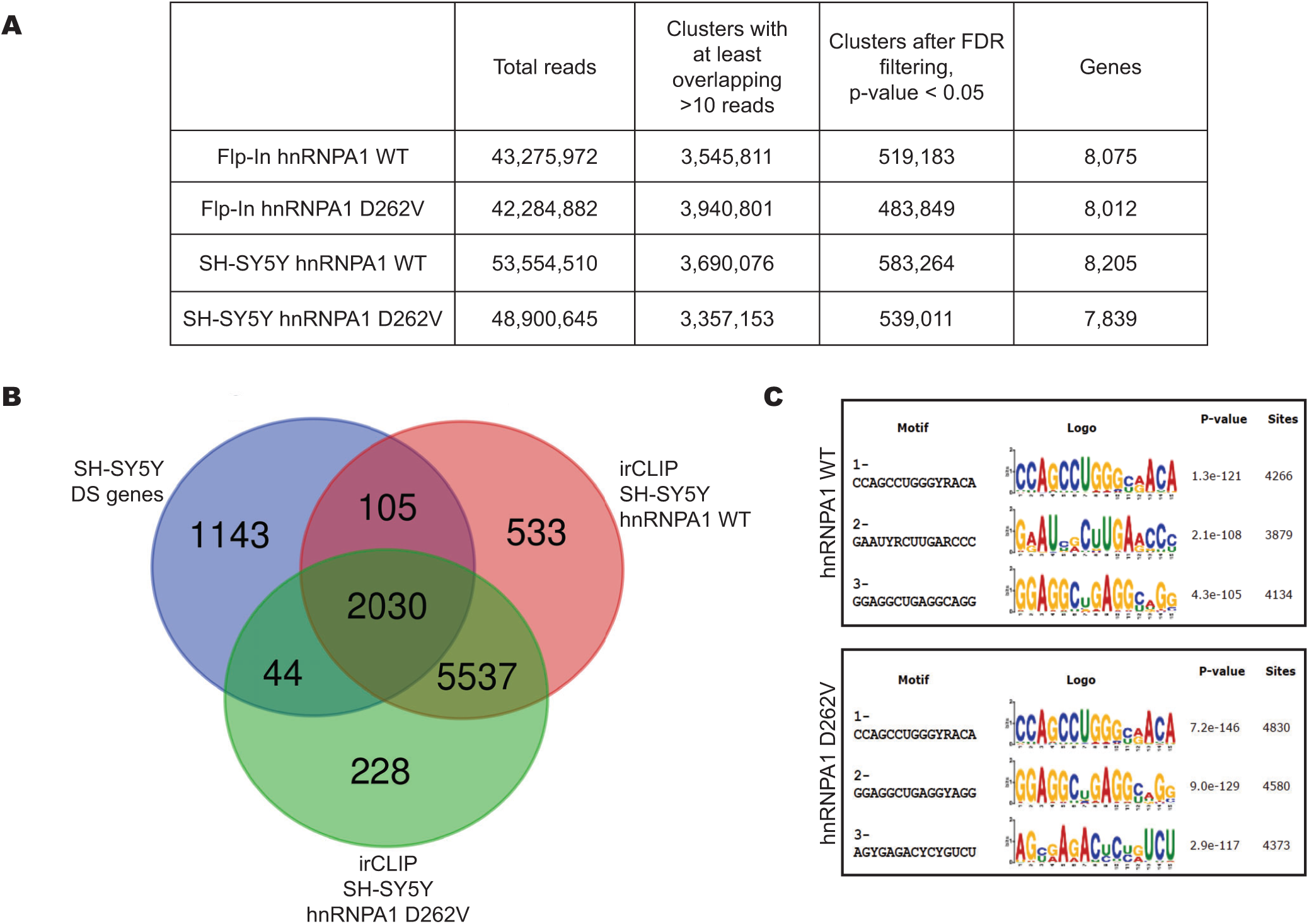
Transcriptome-wide binding maps of hnRNPA1 wild type and D262V mutant. **A**. Summary of irCLIP-seq performed in FLP-In 293 and SH-SY5Y cells expressing hnRNPA1 wild type and D262V mutant proteins. **B**. Venn diagram showing the overlap between the differential splicing target RNAs and transcriptome binding sites of hnRNPA1 wild type and D262V mutant proteins in SH-SY5Y cells. **C**. Motifs derived from hnRNPA1 wild type and D262V binding clusters were generated by extracting k-mers from reads after comparing with control data sets with reads randomly distributed over the same genomic features.

The majority of observed RNA binding targets between the hnRNPA1 wild type and D262V mutant proteins were shared (92-97%), though we did detect 638 transcriptome binding partners that were unique to the hnRNPA1 wild type protein and 272 binding partners unique to the hnRNPA1 D262V mutant protein in SH-SY5Y cells (Fig. 4B). Importantly, comparison of RNA-seq data from SH-SY5Y hnRNPA1 wild type and D262V mutant cells indicates that there are splicing changes associated with differences in hnRNPA1 protein binding. JUM analysis revealed *DNAJC7* gene transcripts are alternative spliced in an intron retention (IR) mode. A recent paper described how exome sequencing in amyotrophic lateral sclerosis implicates a novel gene, DNAJC7, encoding a heat-shock protein (Farhan et al. 2019). It turns out there is an intron retention event [chr17(-):41987910-41988730, ΔPSI =-0.14] observed for transcripts from the DNAJC7 gene in hnRNPA1 mutant-expressing cells. This intron retention event could trigger nonsense-mediated mRNA decay (NMD), hence leading to down-regulation of DNAJC7 protein (Figure 5). While the hnRNPA1 wild type and D262V mutant proteins mostly share common binding sites on the DNAJC7 transcript, only hnRNPA1 wild type binding is observed at the upstream exon junction region (Figure 5, indicated in blue). This lack of exon junction binding by the D262V mutant protein could lead to the intron retention events observed in hnRNPA1 D262V mutant-expressing cells compared to the wild type in SH-SY5Y cells. Several more detailed examples of hnRNPA1 wild type and D262V mutant protein binding to differentially spliced transcripts are shown in the Suppl. Fig. S7-S9. In each case, altered protein binding is correlated with a change in the pre-mRNA splicing pattern.

**Figure 5.**
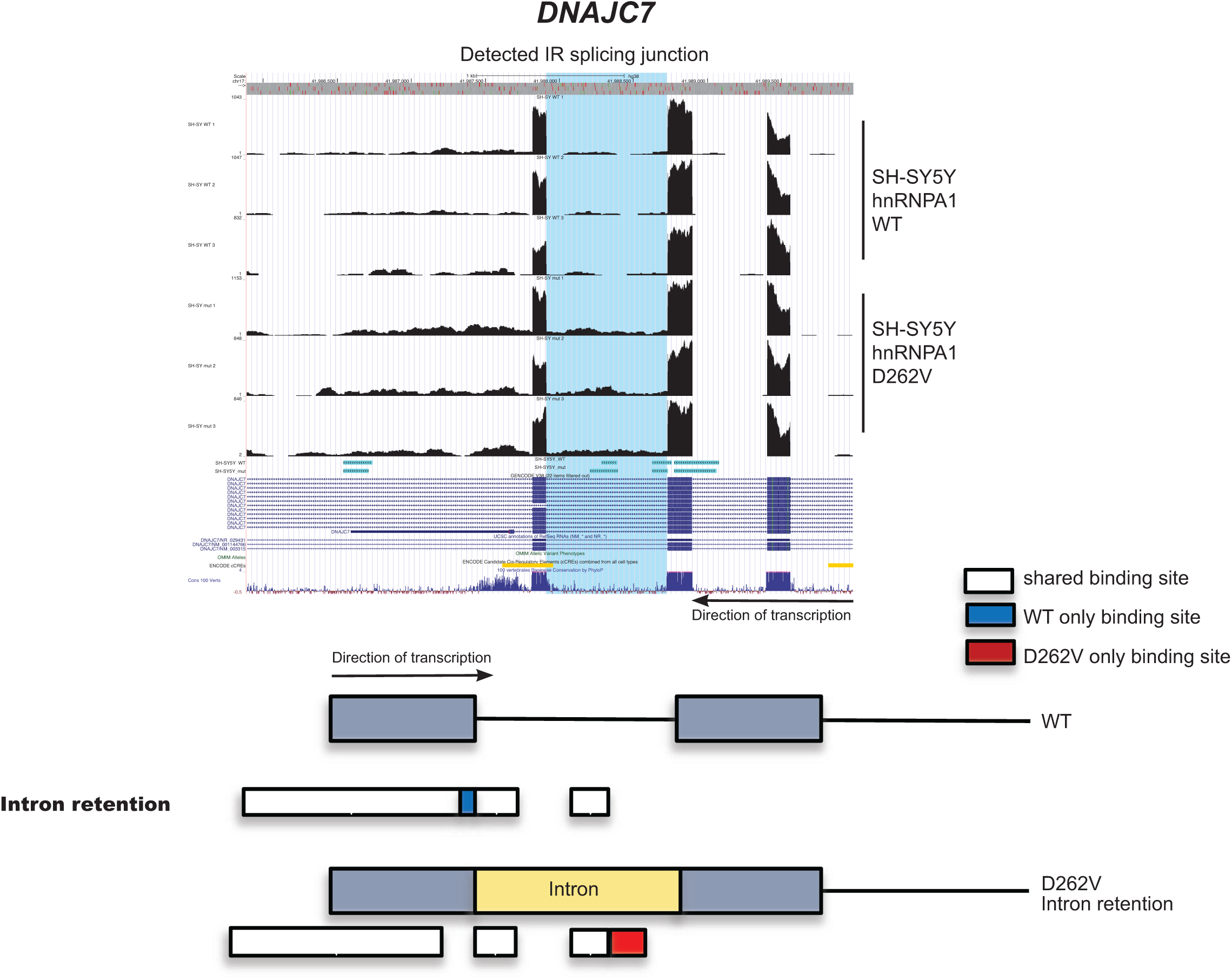
Intron retention in *DNAJC7* transcripts in hnRNPA1 D262V mutant-expressing neuroblastoma cells. *DNAJC* is a newly discovered gene implicated in amyotrophic lateral sclerosis. An intron retention event is observed for the *DNAJC7* transcripts in hnRNPA1 D262V mutant-expressing cells, possibly triggering non-sense-mediated (NMD) mRNA decay that would lead to down-regulation of the *DNAJC7* mRNA and protein.

We also searched our datasets to ask if transcripts encoding known neurodegenerative disease-associated proteins are differentially spliced in cells expressing hnRNPA1 mutant protein (Table 1). These ALS-linked proteins include cytoskeletal proteins, RNA binding proteins, proteins involved in translational control and proteostasis. While GO-term enrichment analysis of DS spliced transcripts mainly yielded DNA repair, cilium organization, response to DNA damage, many known ALS-associated gene transcripts were shown to be differentially spliced in the hnRNPA1 D262V mutant-expressing cell lines, giving more confidence that the splicing targets detected by this RNA-seq analysis are relevant to ALS.

**Table 1.**
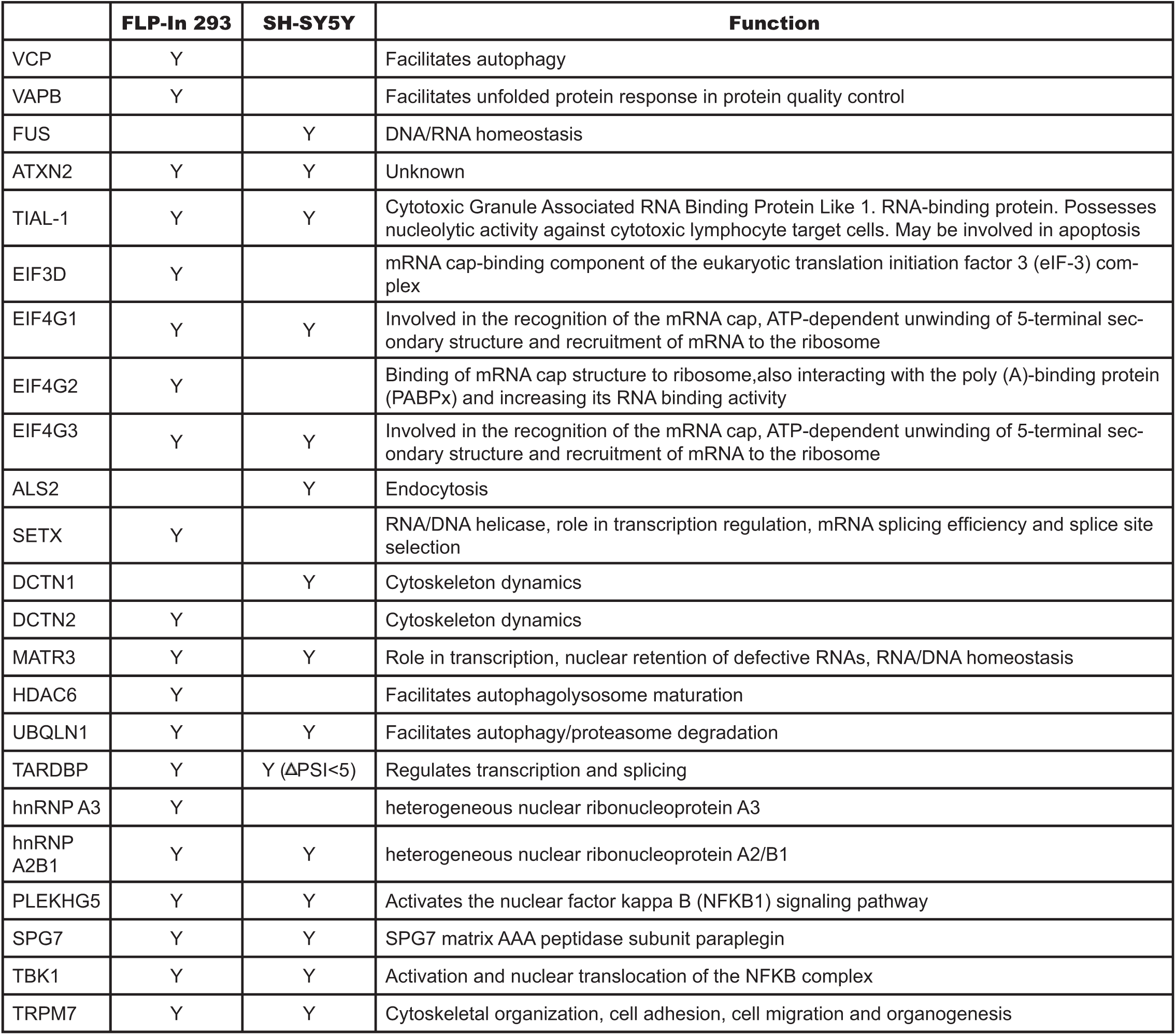
Neurodegenerative disease-associated proteins are identified as high-confidence splicing targets from both SH-SY5Y and in FLP-In™ T-Rex™ 293 hnRNPA1 D262V-expressing cells.

### Proteomic analysis of wild type and D262V mutant proteins

We next wanted to investigate the idea that the hnRNPA1 D262V mutant protein, which exhibits an aggregation phenotype *in vitro* compared to wild type (Kim et al. 2013), might alter protein-protein interactions *in vivo*. Tandem mass tag mass spectrometry (TMT-MS/MS), that enables identification and quantitation of proteins in different samples, was used to make a direct comparison of protein-protein interactions of wild type and mutant hnRNPA1. First, we again optimized immunoprecipitation conditions to completely deplete hnRNPA1 proteins from SH-SY5Y cell extracts expressing either the hnRNPA1 wild type or D262V mutant proteins (Suppl. Fig. S10). Immunoprecipitated proteins were reduced, alkylated, digested with trypsin prior to TMT labeling, and then subjected to LC MS/MS. Our TMT mass spectrometry data indicates that indeed there were differential protein-protein interactions detected between the hnRNPA1 wild type and D262V mutant proteins in cells (Figure 6). Normalized average intensity values were used for quantitation from the TMT-MS/MS results to generate a heat map. We have detected few protein-protein interactions uniquely found only in the hnRNPA1 wild type or D252V mutant samples. Interestingly, we noticed that the hnRNPA1 D262V mutant protein interacted significantly more with other hnRNP proteins, including hnRNPA2B1, hnRNPR, hnRNPH1, hnRNPC, hnRNPL and ribosomal proteins (RPLP0, RPS6, RPS23, RPS25, RPS12, RPS3A, RPLP0P6, RPS15A, RPL4, RPL17, RPL14, RPS17L, RSL1D1, RPL31, and RPS18). It is possible that the hnRNPA1 D262V mutant protein could sequester and possibly trap other RNP-binding proteins or proteins involved in translational control, which could result in altered RNA metabolism or proteostasis changes, both of which are common disease phenotypes of ALS.

**Figure 6.**
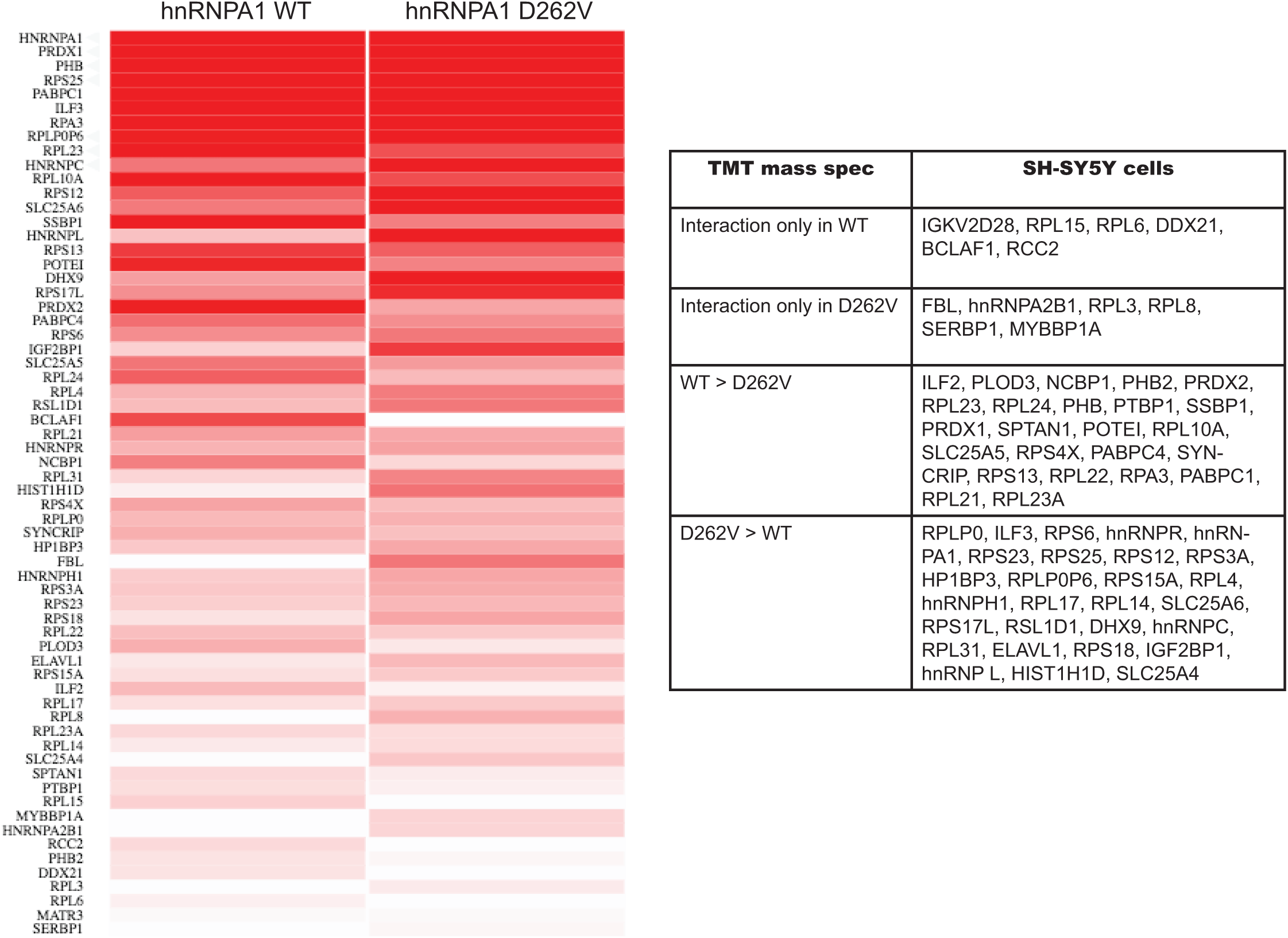
Tandem Mass Tag-Mass Spectrometry (TMT MS) identified differentially interacting proteins in hnRNPA1 wild type and D262V mutant in SH-SY5Y neuroblastoma cells. Heat map (left) of protein abundance differences between hnRNPA1 wild type and D262V mutant immunoprecipitated protein samples. Normalized average ion intensity value quantitation for peptides detected from the indicated proteins using TMT-MS/MS were used for display. A summary of proteins found and/or enriched in different samples is shown on the right.

## Discussion

In order to investigate the mechanisms by which the ALS-linked hnRNPA1 D262V mutation might cause disease, we have taken two different approaches. First, we generated inducible cell lines to express either hnRNPA1 wild type or D262V mutant cDNAs, while expressing endogenous hnRNPA1, using FLP-recombinase integration in 293 cells. Second, we used genome editing with Cas9 at the endogenous hnRNPA1 locus in SH-SY5Y human neuroblastoma cells to generate homozygous hnRNPA1 D262V mutant cell lines. Our studies showed that expression of the hnRNPA1 D262V mutant protein altered pre-mRNA splicing of thousands of transcripts. Gene ontology (GO) term analysis indicated that the genes whose transcripts are differentially spliced are involved in DNA repair, the cellular response to DNA damage stimulus, chromatin remodeling and cilium assembly. We also have shown, using irCLIP *in vivo* binding assays, that the hnRNPA1 D262V mutant protein can either or not bind to some target pre-mRNAs compared to the wild type hnRNPA1 protein. Thus, while the D262V may not affect RNA binding specificity, protein-protein interactions in the C-terminal low complexity domain might affect the affinity of the mutant protein at some binding sites, leading to differential splicing of the target pre-mRNAs. Furthermore, the expression of the mutant hnRNPA1 D262V protein resulted in cells that exhibited a cell aggregation phenotype and a markedly slower cell growth rate. It has been shown that the hnRNPA1 D262V mutant protein, exhibited protein aggregation *in vitro* (Kim et al. 2013; Liu et al. 2016; Beijer et al. 2021). It is possible that the hnRNPA1 D262V mutant protein may sequester and possibly trap other proteins, such as other hnRNPs and/or ribosomal proteins, as evidenced by our TMT-mass spectrometry data. Defects in the function of RNA binding proteins or ribosomes might lead to the observed slow growth phenotype of the hnRNPA1 D262V mutant cells. While the wild type and D262V hnRNPA1 proteins are predominantly nuclear (Suppl Fig.S3 and S4), hnRNPA1 has also been implicated in translational control (Clarke et al. 2021). Thus, we speculate that a single ALS-associated hnRNPA1 D262V mutation encodes a protein that has effects on many aspects of cellular metabolism, including alternative splicing, translation and proteostasis, leading to the altered cell growth and morphology observed here and ultimately leading to the motor neuron death observed in ALS patients.

Interestingly, many neurodegenerative diseases have been associated with defects in primary cilia (Ki et al. 2021). A potential link between autophagy dysfunction and a primary cilia defect has been implicated in Huntington**’**s disease (Kaliszewski et al. 2015). Impaired DNA repair and autophagy dysfunction are two of the proposed mechanisms common in neurodegenerative diseases, along with altered RNA metabolism, impaired protein homeostasis, nucleocytoplasmic transport defects, mitochondrial dysfunction, increased oxidative stress and impaired axonal structure and transport defects. Thus, the splicing defects we have observed in two different cell models are linked to altered mRNA isoforms from genes involved in DNA damage and repair, primary cilia and translation.

Our *in vivo* RNA binding data indicate, as has been previously shown, that hnRNPA1 can bind thousands for RNA transcripts in cells (Huelga et al. 2012; Lee et al. 2018). Our new data indicate that in some cases differential binding of the wild type and D262V mutant hnRNPA1 proteins is correlated with altered splicing patterns of the target transcripts. Previous studies have indicated that hnRNPA1 binding sites can be targeted by anti-sense oligonucleotides (ASOs) in cells and as a treatment for the neurodegenerative disease SMA (Spinal Muscular Atrophy). Identification of hnRNPA1 splicing targets in cells carrying the D262V mutation might also be relevant for transcripts whose splicing is altered in cells carrying other ALS-linked mutations in the RNA binding proteins FUS, TDP-43 and TIA1 (Vance et al. 2009; Liu et al. 2013; Mackenzie et al. 2017). This study provides new information about how the hnRNPA1 D262V mutation might play a role in the development of ALS.

## Materials and Methods

### Cells and generation of hnRNPA1 D262V cell lines

Human SH-SY5Y cells (American Type Culture Collection) were maintained at 37°C in DMEM/F-12 (Gibco™, 11320082) medium supplemented with 10% FBS, 1% Pen-Strep. Wild type and mutant FLAG-tagged hnRNPA1 cDNAs were generated by PCR and cloned into the pcDNA4/FRT/TO. Human FLP-In™ T-REx™ 293 Cell Lines (Invitrogen™, R78007) expressing hnRNPA1 wild type or D262V mutant cDNAs were generated by cotransfecting the hnRNPA1 cDNAs with a FLP recombinase expression plasmid according to the manufacturer’s instructions. Cells were maintained in D-MEM™ (high glucose), 10% FBS, 2mM L-glutamine, and 1% Pen-Strep. 20,000 SH-SY5Y cells were transfected with 6μL (180 pmol) sgRNA (30μM) and 0.5μL of recombinant Cas9 nuclease (20μM). After 10 min. incubation at room temperature, RNP complexes were transfected with 1μL of 100μM mutant donor oligo. A phosphorothioate-modified DNA oligonucleotide (GGATCCTACAATGTTTTTGGC) was used as a donor for Cas9-mediated genome editing. A BamHI restriction site was incorporated into the donor oligonucleotide for screening purposes. After transfection, cells were plated to single cells, grown up and screened by genomic PCR. SH-SY5Y cells had to be plated on irradiated mouse embryo fibroblasts (MEFs) for single cell cloning. Once edited colonies were identified, the presence of the desired mutation was confirmed by Sanger sequencing of the PCR fragments.

### Immunoblotting

Lysates were prepared in RIPA buffer (50mM Tris-HCl, pH 7.4, 150mM NaCl, 1% [v/v] Nonidet P-40, 0.5% [w/v] sodium deoxycholate, 0.1% [w/v] sodium dodecyl sulfate [SDS]) containing protease inhibitors. Equivalent amounts of each sample were subjected to immunoblotting with the following antibodies: hnRNPA1 (Sigma, R4528, clone 9H10, 1:1000 dilution), FLAG (Sigma F3165, 1:1000 dilution), beta actin (Sigma, A2228, 1:3000 dilution).

### Immunofluorescence (IFA) assays

4.5×10^5^ cells were plated 24 hr. before performing IFA. Cells were fixed in 4% paraformaldehyde solution and incubated for 10 min. at 37’C, followed by methanol fixation for 10 min. The cells were block using 6% BSA for 1 hr. at room temperature, followed by incubation with primary antibodies for 2-3 hr. The cells were then washed 3 times in 1X PBS/0.05% Tween 20, then incubated with secondary antibodies followed by PBS/0.05% Tween washes. IFA was performed using hnRNPA1 (Sigma, R4528; 1:500 dilution), eIF3α (Abcam, 3411S; 1:500 dilution) and Alexa Fluor 488- or 568-labeled donkey anti-rabbit or goat anti-mouse secondary antibodies (1:1,500 dilution, Life Technologies, A10042, A11029), and mounted with DAPI-containing VECTASHIELD mounting medium (Vector Labs). The images were obtained at the CNR Biological Imaging Facility, University of California at Berkeley.

### RNA-seq library preparation and sequencing

RNA isolation was performed using the RNeasy Minikit (Qiagen, 74104), followed by 30min. of DNase treatment (Ambion, AM2238) at 37°C, poly(A)^+^ RNA transcripts were isolated [NEBNext poly(A) mRNA magnetic isolation module; New England Biolabs, E7490] from 1μg of total RNA for RNA library preparation and sequencing using NEBNext Ultra Directional RNA Library Preparation Kit for Illumina (New England Biolabs, E7420S) according to the manufacturer’s instruction. The samples were sequenced on an Illumina HiSeq 4000 (FLP-In 293) or NovaSeq SP (SH-SY5Y) with 100-bp paired-end reads at the Vincent J. Coates Genomics Sequencing Laboratory at the University of California at Berkeley.

### Analysis of pre-mRNA splicing patterns using JUM and differential expression analysis using DESeq2

Pre-mRNA splicing analysis using JUM.2.0.2 software to detect pre-mRNA splicing patterns was performed as described before (Lee et al. 2018). A *P*-value of 0.05 was used as the statistical cutoff for differentially spliced AS events. Expressed genes for GO-terminology enrichment analysis was generated using cufflinks.v2.1.1(Trapnell et al. 2009) using the following parameters: --library-type fr-firststrand -m 100 -s 10, then filtered for FPKM > 1.

For Differential expression analysis, uniquely mapped reads were mapped to hg38 using STAR_2.5.3a_modified version using --outFilterMultimapNmax 1. Read counts were calculated using HTSeq script htseq-count with the following specifications: -s yes -r pos -f. Differential gene expression analysis was performed using DESeq2 (Love et al. 2014) using CPM cutoff.

The expression cutoff (CPM) value was determined according to the library size and normalization factors, roughly equal to 10 */L* where *L* is the minimum library size in millions (Chen Y 2016).

### Quick-irCLIP

To identify the RNAs bound by hnRNPA1 wild type versus D262V mutant, we performed quick-irCLIP, cross-linking and inferred immunoprecipitation, according to the previously published protocol (Kaczynski et al. 2019), with minor modifications. Library was prepared using SMARTer smRNA-seq Kit for Illumina (Takara, 635029) according to the manufacturer’s instructions. The samples were sequenced on an Illumina NovaSeq S1 with 100-bp paired-end reads at the Vincent J. Coates Genomics Sequencing Laboratory at the University of California at Berkeley.

The Quick irCLIP-seq reads were preprocessed prior to mapping using FASTX-Toolkit (http://hannonlab.cshl.edu/fastx_toolkit). Reads were quality-filtered based on quality score (fastq_quality_filter -q 25 -p 80), and PCR duplicates were collapsed (fastx_collapser). Only uniquely mapped reads were used for further analysis. The reads were mapped to hg38 using STAR using the following parameters: --outFilterMultimapNmax 1 --quantMode TranscriptomeSAM GeneCounts --outReadsUnmapped fastx --outSAMtype BAM SortedByCoordinate. Uniquely mapped reads were filtered and then used for cluster identification using pycrac/1.2.3.0 (Webb et al. 2014). irCLIP clusters were generated using at least ten overlapping unique cDNA reads generated after removal of PCR duplicates (pyClusterReads.py --cic=10). pyCalculateFDRs were used to filter out statistically significant clusters over the regions with a read coverage of at least 5 (--min=10) and FDR p < 0.05. De novo binding motifs for hnRNPA1 wild type and mutant protein were determined using STREME (Bailey 2021) using the following parameter: streme --verbosity 1 --oc --rna --minw 8 --maxw 15 --pvt 0.05.

### Tandem mass tag (TMT) mass spectrometry

The protocol we used was modified slightly from a previous study (Griesser et al. 2020). 500μg cell lysate from SH-SY5Y cells was incubated with 50μg hnRNPA1 antibody (Sigma R4528) bound to Dynabeads™ Protein G beads in lysis buffer (0.5% n-dodecyl-b-D-maltoside, 150mM NaCl, 50mM HEPES-NaOH, pH 7.4). The beads were washed in 4X in 0.1%/Triton-X 100 in 25mM HEPES buffer (pH 7.4). The samples were treated with 10mM DTT to reduce reversibly oxidized cysteines for 45 min. at 50°C and then were alkylated with 20mM iodoacetic acid for 45 min. at room temperature with shaking at 1000 rpm. The samples were subjected to SP3 digestion and TMT labeled exactly as previously described (Griesser et al. 2020). The mass spectrometry was performed at the Vincent J. Coates Proteomics/Mass Spectrometry Laboratory, UC Berkeley. Normalized average intensity values quantitation from TMT-MS/MS were visualized using Clustergrammer (Fernandez et al. 2017).

## Data deposition

The data reported will be deposited in the Gene Expression Omnibus (GEO) database upon the acceptance of the manuscript.

## Acknowledgments

We thank George Ghanim (current address: MRC Laboratory of Molecular Biology, Cambridge, UK) for designing of hnRNPA1 donor oligonucleotide. We thank Dirk Hockemeyer and his lab for providing irradiated mouse embryo fibroblasts (MEFs) to use as feeder cells for the SH-SY5Y cells. This work used the Vincent J. Coates Genomics Sequencing Laboratory at University of California at Berkeley, supported by National Institutes of Health (NIH) S10 Instrumentation grants S10RR025622, S10RR029668, and S10RR027303. This research used the Savio computational cluster resource provided by the Berkeley Research Computing program at the University of California at Berkeley (supported by the University of California at Berkeley Chancellor, Vice Chancellor for Research, and Chief Information Officer). IFA images were taken at the CNR Biological Imaging Facility at the University of California, Berkeley, which was supported in part by the National Institutes of Health S10 program under award number 1S10RR026866-01. This work used the Vincent J. Proteomics/Mass Spectrometry Laboratory at UC Berkeley, supported in part by NIH S10 Instrumentation Grant S10RR025622. We thank Lori Kohlstaedt of the Vincent J. Proteomics/Mass Spectrometry Laboratory at UC Berkeley for assistance in the analysis of mass spectrometry data. This work was funded by the NIH grants R01 GM097352 and R35 GM118121, awarded to D.C.R.

## Author Contributions

D.C.R. and Y.J.L. conceived the study. Y.J.L. performed experiments, analyzed data and generated figures. Y.J.L. and D.C.R. wrote the original draft. Y.J.L. and D.C.R. reviewed and edited the manuscript. D.C.R. obtained funding.

## Figure Legends

**Supplemental Figure 1S. Differential expression analysis comparing FLP-In 293 hnRNPA1 D262V to the wildtype**. A volcano plot of the differential expression statistics showing genes that are down-regulated (in red) or up-regulated (in blue) with a log fold change (FC) > 1.5 and an FDR-adjusted p-value < 0.05. A total of 345 genes were differentially expressed in FLP-In 293 hnRNPA1 D262V cells.

**Supplemental Figure 2S. Differential expression analysis comparing SH-SY5Y hnRNPA1 D262V to the wildtype**. A volcano plot of the differential expression statistics showing genes that are down-regulated (in red) or up-regulated (in blue) with a log fold change (FC) > 1.5 and an FDR-adjusted p-value < 0.05. 619 genes are differentially expressed.

**Supplemental Figure S3: hnRNPA1 D262V mutant cells in FLP-In™ T-Rex™ 293 cells and SH-SY5Y human neuroblastoma cells exhibit distinct cell phenotype, including cell aggregation and slow growth rate. A**. Tetracycline (Tet)-inducible 293 FLP-In hnRNPA1 wild type and hnRNPA1 mutant (D262V) cells were induced with 1μg/mL Tetracycline (Tet) for 72 hr. prior to immunofluorescence assay using anti-FLAG to detect exogenously expressed hnRNPA1. **B**. Growth curve of 293 FLP-In hnRNPA1 WT and hnRNPA1 mutant (D262V). Cells were treated with 1μg/mL tetracycline (Tet) for the indicated number of days and cell numbers were counted. **C**. SH-SY5Y cells were cultured for 10 days after plating 1×10^5^ cells/well preceding crystal violet staining. **D**. Differential Interference Contrast (DIC) imaging on a confocal microscope.

**Supplemental Figure S4**. Delayed stress-granule (SG) disassembly in hnRNPA1 D262V neuroblastoma cells. SH-SY5Y neuronal cell line expressing hnRNPA1 wild type or D262V mutant were treated with 500 μM arsenite treatment for 30 min., then followed SG assembly by immunofluorescence assay using eIF3α antibody which has been shown to translocate to SG under stress. hnRNPA1 is indicated in green and eIF3α is indicated in red.

**Supplemental Figure S5**. Differentially spliced transcripts in FLP-In 293 hnRNPA1 D262V and SH-SY5Y hnRNPA1 D262V neuroblastoma cells include neuronal RNP granule-associated RNAs, eIF4A associated RNAs and stress-granule (SG) associated RNAs.

**Supplemental Figure S6. Quick-irCLIP in FLP-In 293 cells expressing hnRNPA1 WT and D262V mutant. A**. Tetracycline(Tet)-inducible 293 FLP-In hnRNPA1 WT and hnRNPA1 mutant (D262V) cells were induced with 1μg/mL Tetracycline (Tet) for 72 hr. prior to harvest, followed by nuclear extract preparation. Efficient pulldown of FLAG-tagged exogenous hnRNPA1 wild type and D262V shown in immunoblot analysis. **B**. Odyssey CLx infrared scans of the nitrocellulose bound immunoprecipitates using FLAG-antibody against exogenously expressed hnRNPA1wild type and D262V mutant in FLP-In 293 cells.

**Supplemental Figure S7. Expression of hnRNPA1 D262V mutant leads of exon 8 inclusion of hnRNPA1 mRNA that results in isoform change**. Genome browser shot [chr12(+):54282826-54284256] of the hnRNPA1 gene with RNA-seq data upon expression of hnRNPA1 wild type (top) and D262V (bottom). hnRNPA1 wild type-specific binding site is indicated in blue box, while hnRNPA1 D262V-specific binding site is indicated in red. Shared binding sites are indicated in white. Expression of hnRNPA1 D262V mutant in SH-SY5Y cells leads to inclusion of exon 8 that leads to an isoform switch.

**Supplemental Figure S8. Expression of hnRNPA1 D262V mutant leads of exon inclusion of ZNF207 transcript**. ZNF207 is a kinetochore- and microtubule-binding protein that plays a key role in spindle assembly. Genome browser shot of the hnRNPA1 gene with RNA-seq data upon expression of hnRNPA1 wild type (top) and D262V (bottom). On hnRNPA1 wild type-specific binding site is observed nearby cassette exon region [chr17(+):32365487-32367770], indicated in blue box. Expression of hnRNPA1 D262V mutant in SH-SY5Y cells leads to increase in exon inclusion compared to the unedited control.

**Supplemental Figure S9. Distal A3SS site usage of hnRNPA1 D262V mutant on POGZ transcript**. POGZ (Pogo Transposable Element Derived With ZNF Domain) plays a role in mitotic cell cycle progression and is involved in kinetochore assembly and mitotic sister chromatid cohesion. Expression of hnRNPA1 D262V leads to usage of distal A3SS [chr1(-):151428041-151428121] compared to proximal A3SS usage in hnRNPA1 wild type-expressing cells. hnRNPA1 wild type-specific binding site is indicated in the blue box.

**Supplemental Figure S10**. Efficient pulldown of hnRNPA1 wild type and D262V proteins in SH-SY5Y cells shown by immunoblot analysis. Immunoprecipitated proteins were reduced, alkylated, digested with trypsin prior to TMT labeling, and then subjected to LC MS/MS.

