## Supplemental Figures for "Analysis of altered pre-mRNA splicing patterns caused by a mutation in the RNA binding protein hnRNPA1 linked to amyotrophic lateral sclerosis"

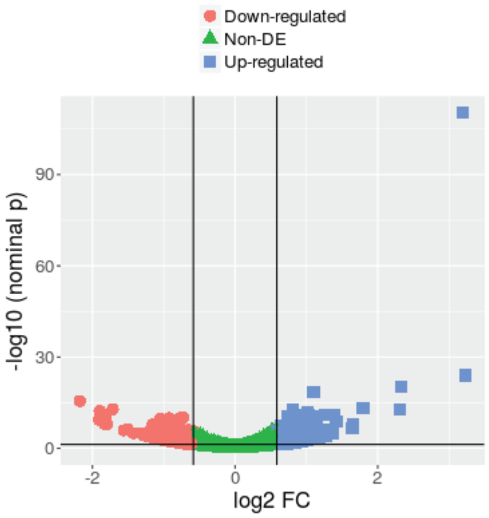

FDR p-value < 0.05, FC > 1.5

| Genes | FDR < 0.05, FC > 1.5 |
|---|---|
| Down-regulated | 155 |
| Up-regulated | 190 |

| GO term | Description | P-value |
|---|---|---|
| GO:0046098 | guanine metabolic process | 3E-4 |
| GO:0045047 | protein targeting to ER | 3.65E-4 |
| GO:0072599 | establishment of protein localization to endoplasmic reticulum | 4.75E-4 |
| GO:0072657 | protein localization to membrane | 5.53E-4 |
| GO:0006614 | SRP-dependent cotranslational protein targeting to membrane | 7.23E-4 |
| GO:0046037 | GMP metabolic process | 7.68E-4 |
| GO:0070972 | protein localization to endoplasmic reticulum | 9.73E-4 |

**Supplemental Figure 1S. Differential expression analysis comparing FLP-In 293 hnRNPA1 D262V to the wildtype.** A volcano plot of the differential expression statistics showing genes that are down-regulated (in red) or up-regulated (in blue) with a log fold change (FC) > 1.5 and an FDR-adjusted p-value < 0.05. A total of 345 genes were differentially expressed in FLP-In 293 hnRNPA1 D262V cells.

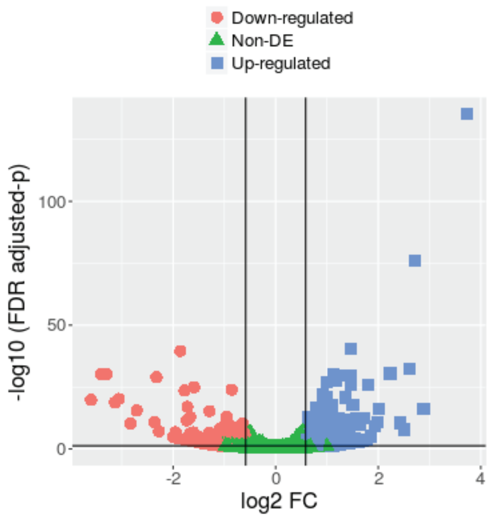


FDR p-value < 0.05, FC > 1.5

| Genes | FDR < 0.05, FC > 1.5 |
|---|---|
| Down-regulated | 267 |
| Up-regulated | 352 |

| GO term | Description | P-value |
|---|---|---|
| GO:0051642 | centrosome localization | 8.82E-5 |
| GO:0061842 | microtubule organizing center localization | 1.12E-4 |
| GO:0035195 | gene silencing by miRNA | 5.73E-4 |
| GO:0051620 | norepinephrine uptake | 9.46E-4 |

**Supplemental Figure 2S. Differential expression analysis comparing SH-SY5Y hnRNPA1 D262V to the wildtype.** A volcano plot of the differential expression statistics showing genes that are down-regulated (in red) or up-regulated (in blue) with a log fold change (FC) > 1.5 and an FDR-adjusted p-value < 0.05. 619 genes are differentially expressed.

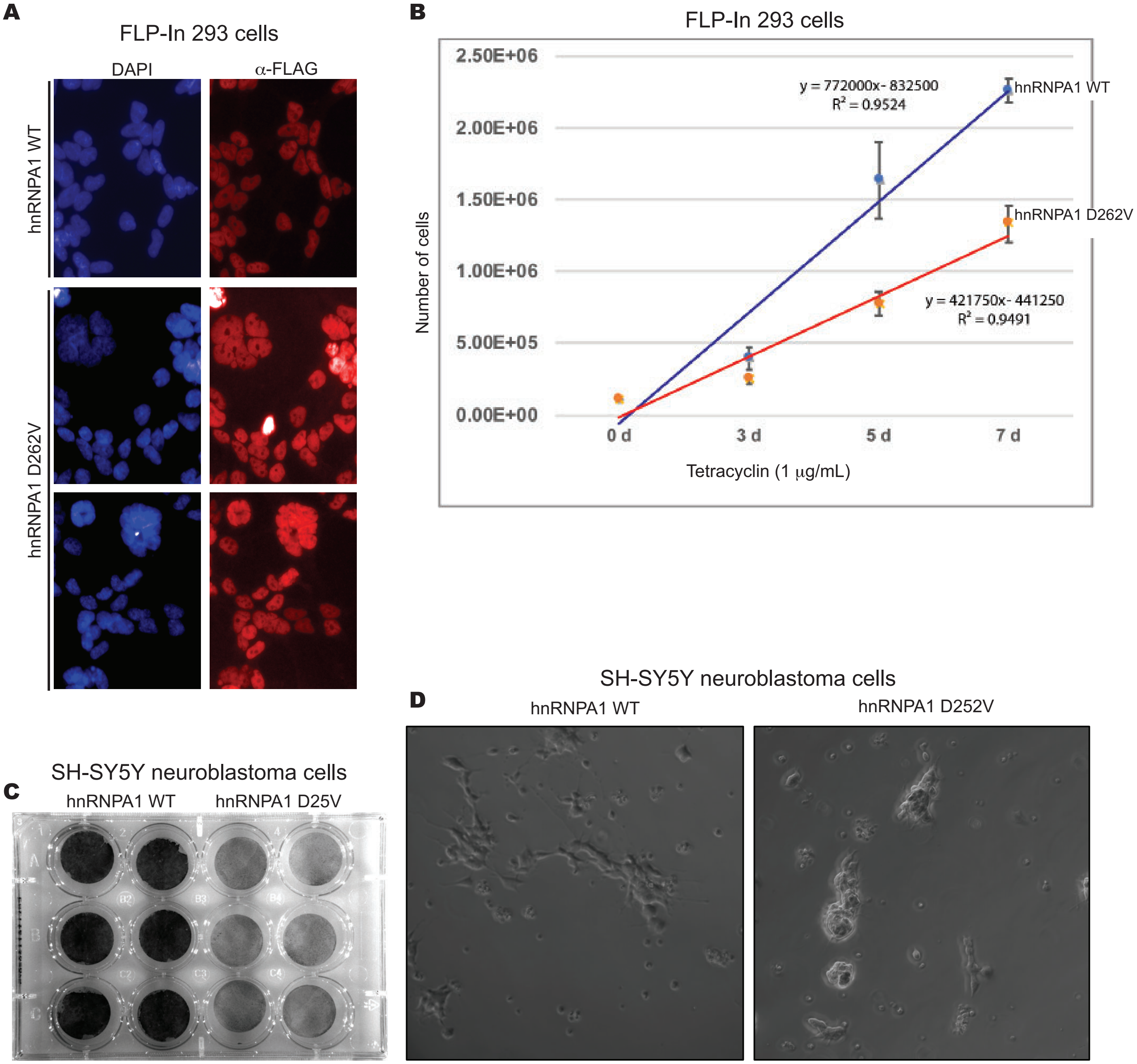


**Supplemental Figure S3: hnRNPA1 D262V mutant cells in FLP-In™ T-Rex™ 293 cells and SH-SY5Y human neuroblastoma cells exhibit distinct cell phenotype, including cell aggregation and slow growth rate.** A. Tetracycline (Tet)-inducible 293 FLP-In hnRNPA1 wild type and hnRNPA1 mutant (D262V) cells were induced with 1µg/mL Tetracycline (Tet) for 72 hr. prior to immunofluorescence assay using anti-FLAG to detect exogenously expressed hnRNPA1. B. Growth curve of 293 FLP-In hnRNPA1 WT and hnRNPA1 mutant (D262V). Cells were treated with 1µg/mL tetracycline (Tet) for the indicated number of days and cell numbers were counted. C. SH-SY5Y cells were cultured for 10 days after plating 1x105 cells/well preceding crystal violet staining. D. Differential Interference Contrast (DIC) imaging on a confocal microscope.

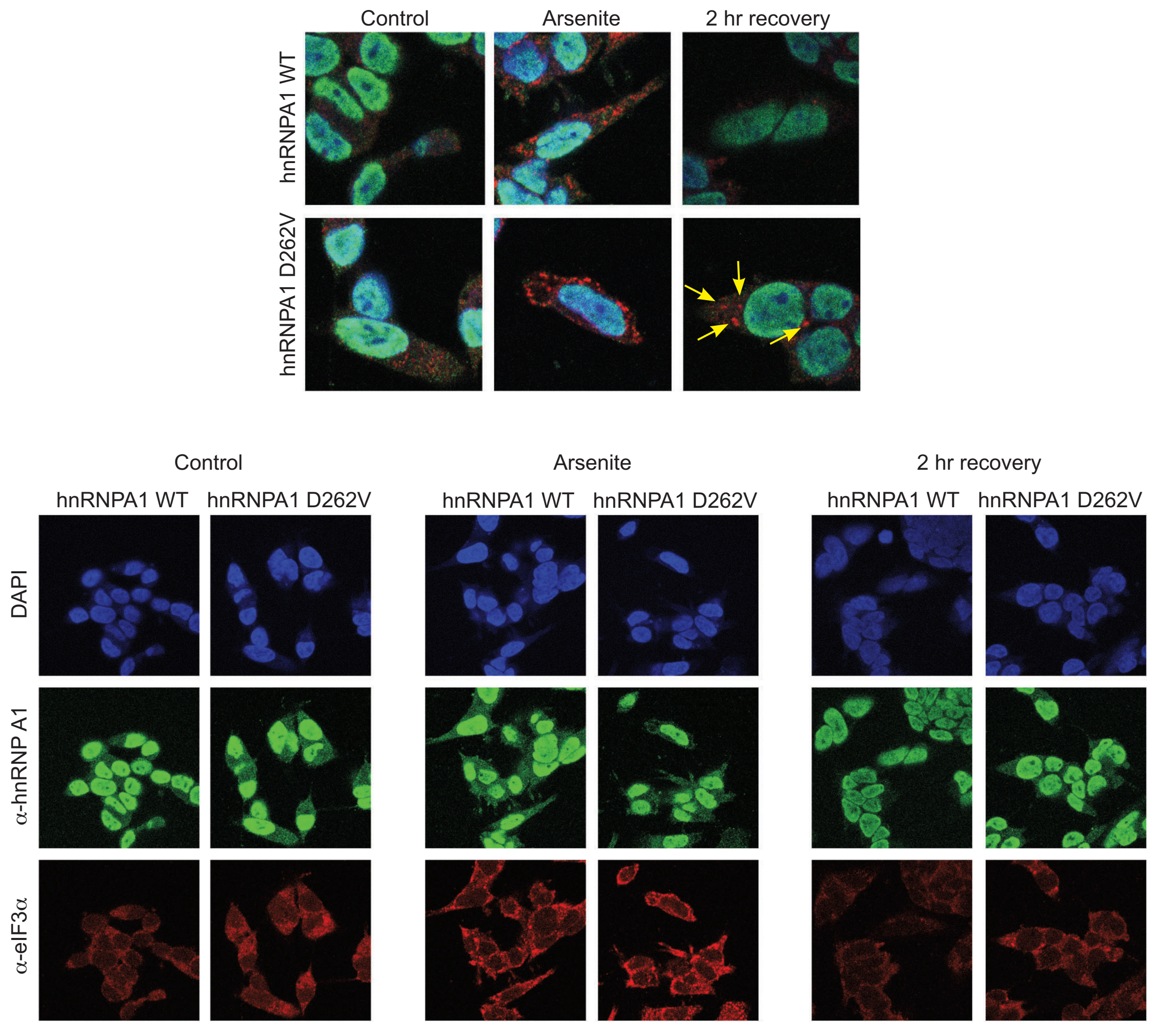


**Supplemental Figure S4. Delayed stress-granule (SG) disassembly in hnRNPA1 D262V neuroblastoma cells.** SH-SY5Y neuronal cell line expressing hnRNPA1 wild type or D262V mutant were treated with 500 μM arsenite treatment for 30 min., then followed SG assembly by immunofluorescence assay using eIF3α antibody which has been shown to translocate to SG under stress. hnRNPA1 is indicated in green and eIF3α is indicated in red.

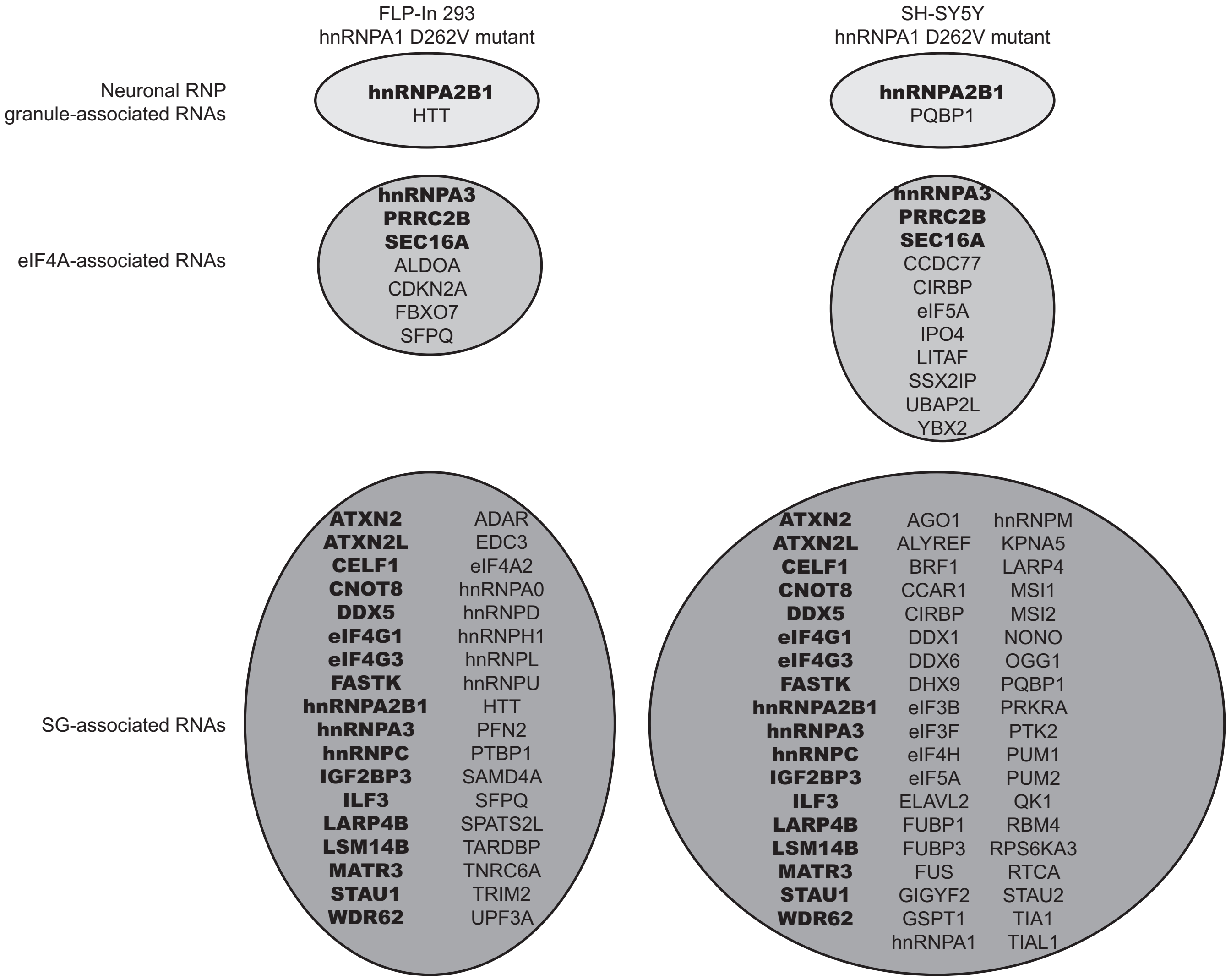


**Supplemental Figure S5.** Differentially spliced transcripts in FLP-In 293 hnRNPA1 D262V and SH-SY5Y hnRNPA1 D262V neuroblastoma cells include neuronal RNP granule-associated RNAs, eIF4A associated RNAs and stress-granule (SG) associated RNAs.

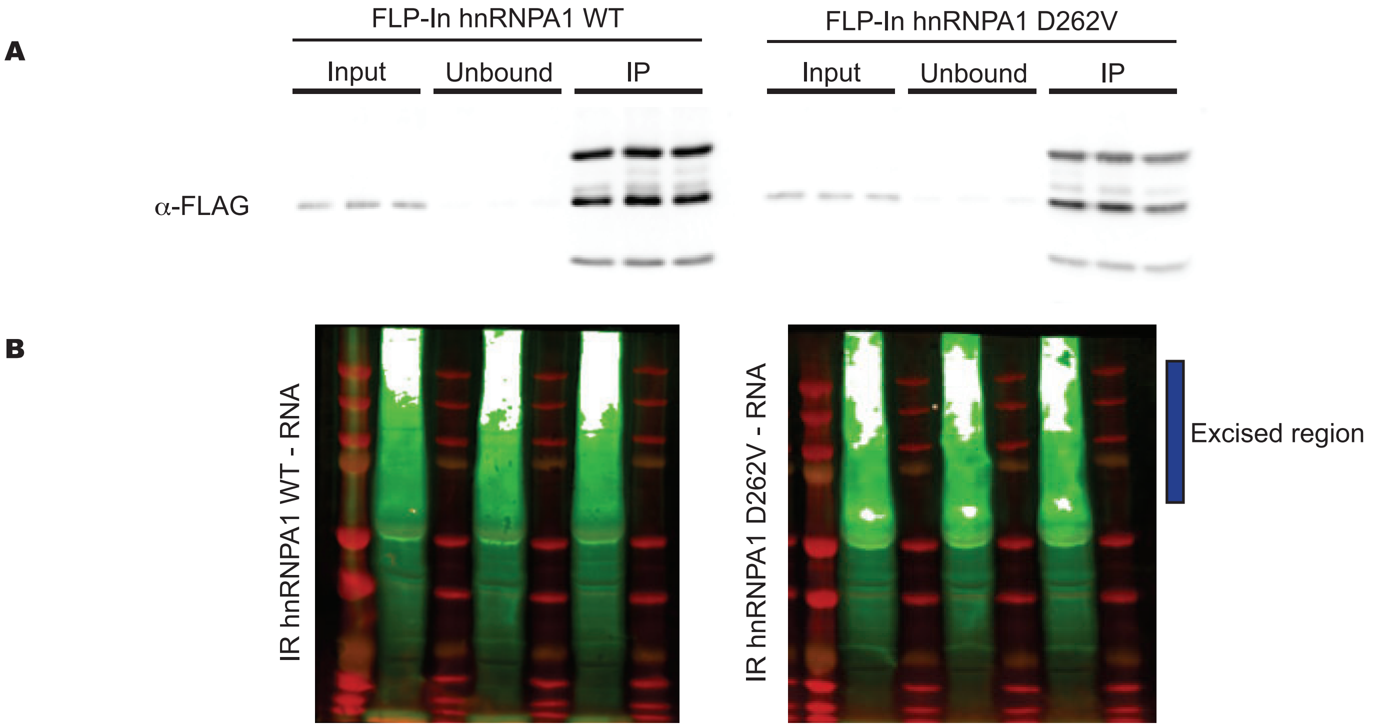


**Supplemental Figure S6. Quick-irCLIP in FLP-In 293 cells expressing hnRNPA1 WT and D262V mutant.** A. Tetracycline(Tet)-inducible 293 FLP-In hnRNPA1 WT and hnRNPA1 mutant (D262V) cells were induced with 1μg/mL Tetracycline (Tet) for 72 hr. prior to harvest, followed by nuclear extract preparation. Efficient pulldown of FLAG-tagged exogenous hnRNPA1 wild type and D262V shown in immunoblot analysis. B. Odyssey CLx infrared scans of the nitrocellulose bound immunoprecipitates using FLAG-antibody against exogenously expressed hnRNPA1wild type and D262V mutant in FLP-In 293 cells.

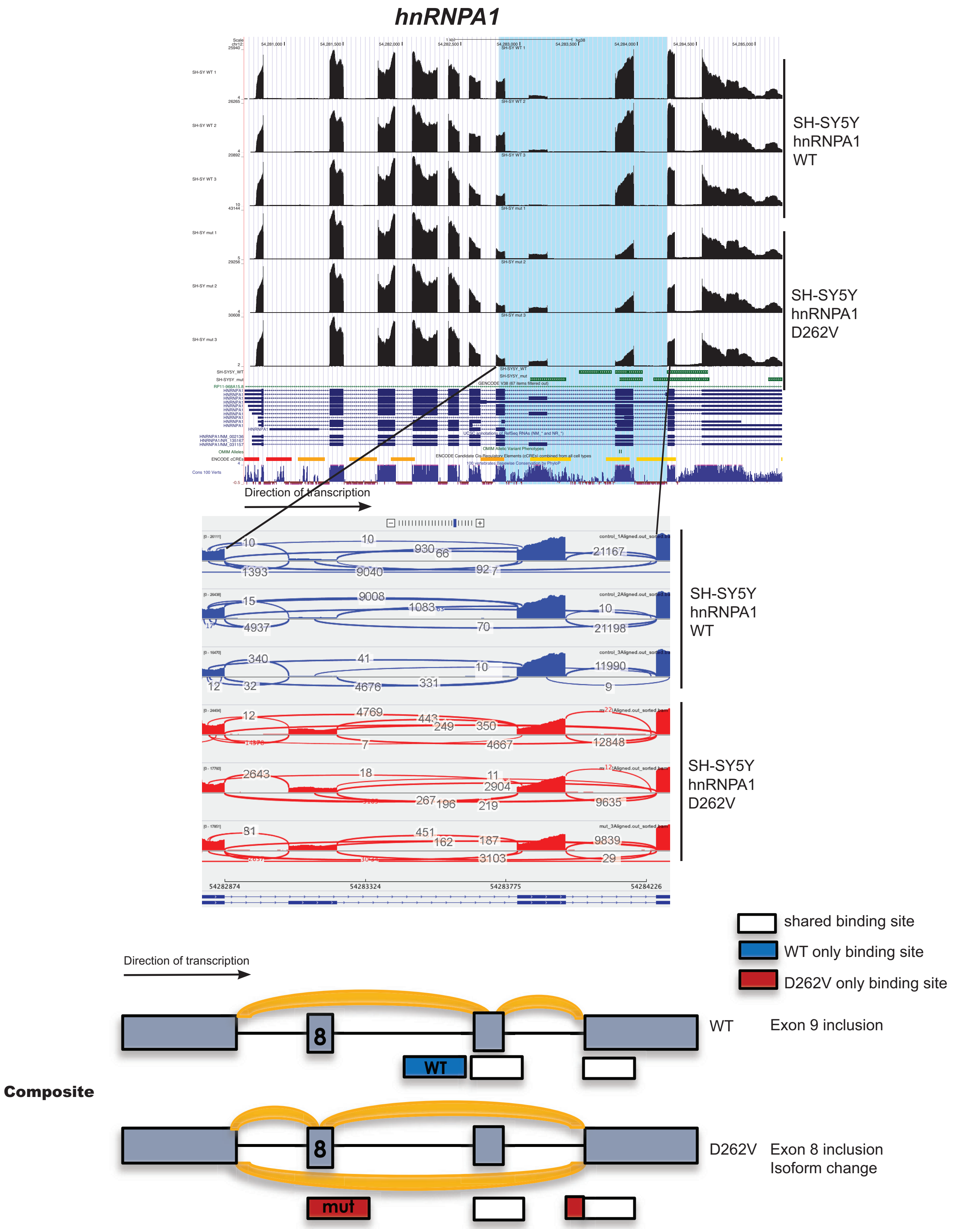


**Supplemental Figure S7. Expression of hnRNPA1 D262V mutant leads of exon 8 inclusion of hnRNPA1 mRNA that results in isoform change.** Genome browser shot [chr12(+):54282826-54284256] of the hnRNPA1 gene with RNA-seq data upon expression of hnRNPA1 wild type (top) and D262V (bottom). hnRNPA1 wild type-specific binding site is indicated in blue box, while hnRNPA1 D262V-specific binding site is indicated in red. Shared binding sites are indicated in white. Expression of hnRNPA1 D262V mutant in SH-SY5Y cells leads to inclusion of exon 8 that leads to an isoform switch.

### *ZNF207*

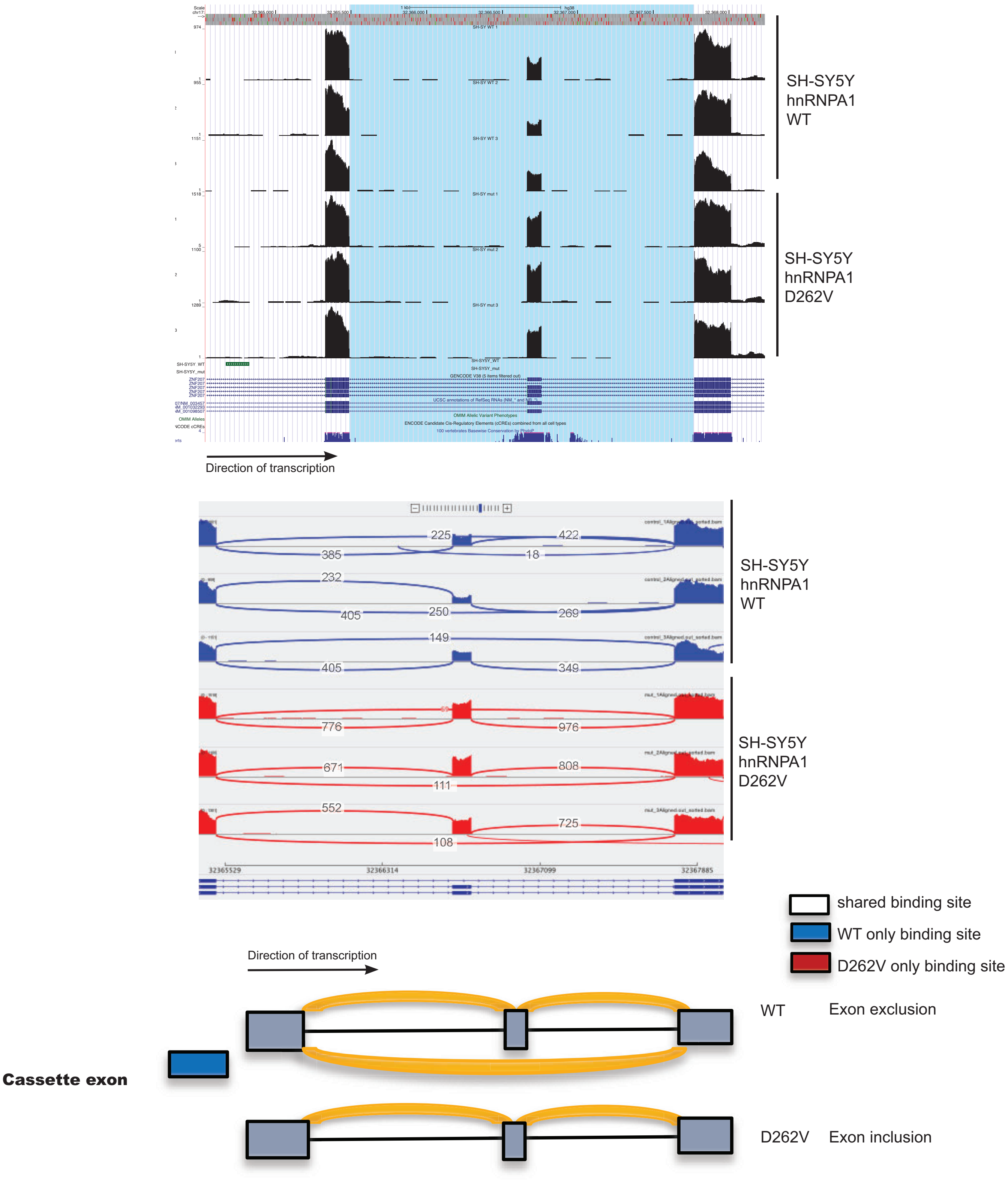


**Supplemental Figure S8. Expression of hnRNPA1 D262V mutant leads of exon inclusion of ZNF207 transcript.** ZNF207 is a kinetochore- and microtubule-binding protein that plays a key role in spindle assembly. Genome browser shot of the hnRNPA1 gene with RNA-seq data upon expression of hnRNPA1 wild type (top) and D262V (bottom). On hnRNPA1 wild type-specific binding site is observed nearby cassette exon region [chr17(+):32365487-32367770], indicated in blue box. Expression of hnRNPA1 D262V mutant in SH-SY5Y cells leads to increase in exon inclusion compared to the unedited control.

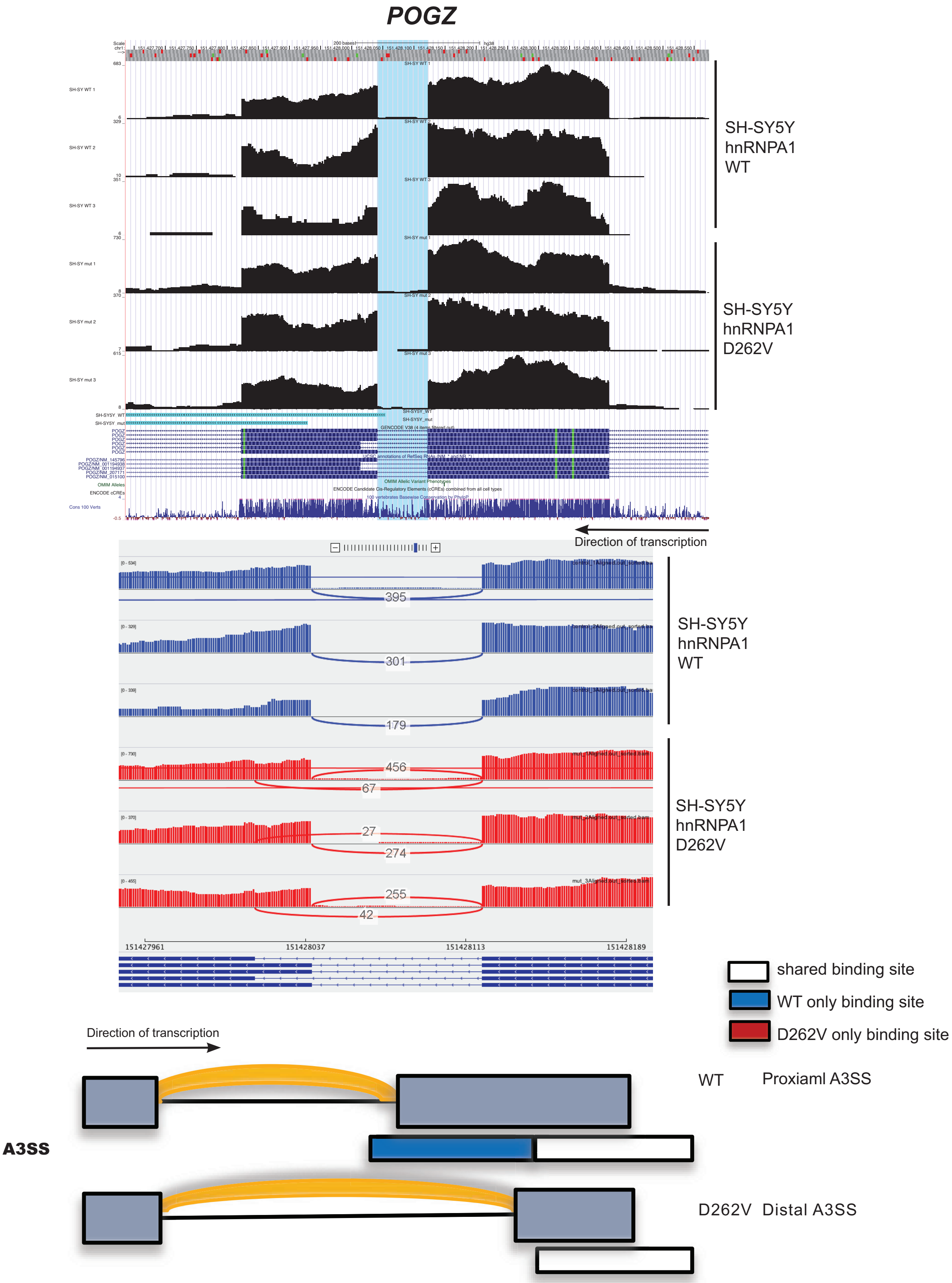


**Supplemental Figure S9. Distal A3SS site usage of hnRNPA1 D262V mutant on POGZ transcript.** POGZ (Pogo Transposable Element Derived With ZNF Domain) plays a role in mitotic cell cycle progression and is involved in kinetochore assembly and mitotic sister chromatid co-hesion. Expression of hnRNPA1 D262V leads to usage of distal A3SS [chr1(-):151428041-151428121] compared to proximal A3SS usage in hnRNPA1 wild type-expressing cells. hnRNPA1 wild type-specific binding site is indicated in the blue box.

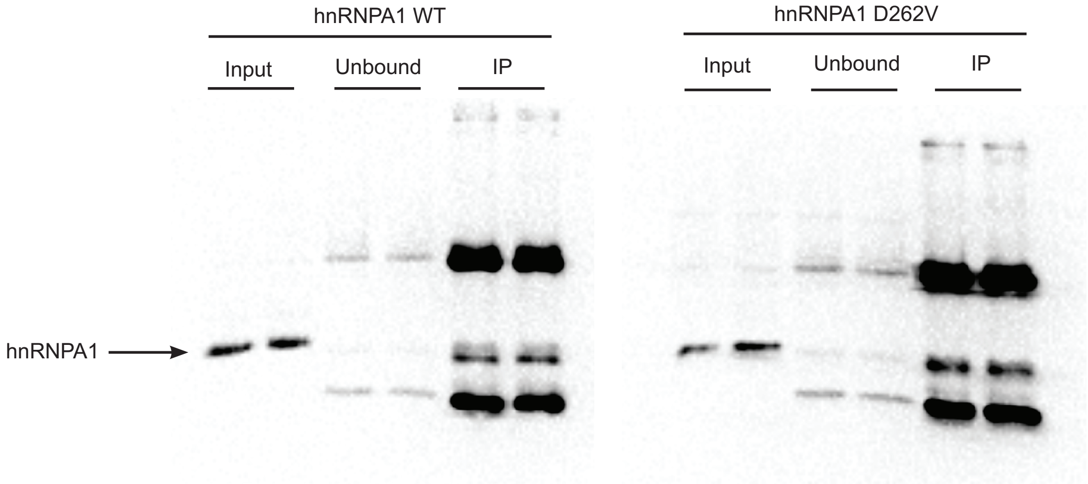


**Supplemental Figure S10. Efficient pulldown of hnRNPA1 wild type and D262V proteins in SH-SY5Y cells shown by immunoblot analysis.** Immunoprecipitated proteins were reduced, alkylated, digested with trypsin prior to TMT labeling, and then subjected to LC MS/MS.
